# Phase-amplitude coupling of NREM sleep oscillations is unaffected by pre-sleep learning but related to overnight memory gains depending on the declarative learning paradigm

**DOI:** 10.1101/2023.11.23.568501

**Authors:** Nathan Cross, Jordan O’Byrne, Oren M. Weiner, Julia Giraud, Aurore A. Perrault, Thien Thanh Dang-Vu

## Abstract

There is growing evidence in humans linking the temporal coupling between spindles and slow oscillations during NREM sleep with the overnight stabilization of memories encoded from daytime experiences in humans. However, whether the type and strength of learning influence that relationship is still unknown. Here we tested whether the amount or type of verbal word-pair learning prior to sleep affects subsequent phase-amplitude coupling (PAC) between spindles and slow oscillations (SO). We measured the strength and preferred timing of such coupling in the EEG of 41 healthy human participants over a post-learning and control night, to compare intra-individual changes with inter-individual differences. We leveraged learning paradigms of varying word-pair (WP) load: 40 WP learned to a minimum criterion of 60% correct (n=11); 40 WP presented twice (n=15); 120 WP presented twice (n=15). There were no significant differences in the preferred phase or strength between the control and post-learning nights, in all learning conditions. We observed an overnight consolidation effect (improved performance at delayed recall) for the criterion learning condition only, and only in this condition was the overnight change in memory performance significantly positively correlated with the phase of SO-spindle coupling. These results suggest that the coupling of brain oscillations during human NREM sleep are stable traits that are not modulated by the amount of pre-sleep learning, yet are implicated in the sleep-dependent consolidation of memory.

## Introduction

Sleep contributes significantly to memory consolidation, however, the mechanisms underlying sleep-dependent memory consolidation are still not well established. The active consolidation hypothesis predicts that sleep is a brain state optimized for active redistribution of temporary memory representations into long-term stores (Born and Wilhelm, 2012). This hypothesis asserts that the process of consolidation relies on the repeated reactivation of the newly encoded memories during off-line periods (Born and Wilhelm, 2012; Goto and Hayashi, 2023). Such off-line periods occur regularly during sleep, in fact the neocortex ‘reboots’ itself from complete silence of neuronal activity (“down” states) hundreds to thousands of times during non-REM (NREM) sleep (Buzsáki, 2019; Steriade et al., 1993). These events have been termed slow oscillations and are a major hallmark feature of NREM sleep (Neske, 2016).

The reactivation of memories during NREM sleep are proposed to be regulated by an interchange between the hippocampus and neocortex, which is effectively under feed-forward control of the slow oscillation (Born and Wilhelm, 2012; Steriade, 2006). Cross-frequency phase-amplitude coupling (PAC) analyses have revealed a form of hierarchical nesting of brain oscillations, such that cortical slow oscillations modulate the timing of sleep spindles generated by thalamocortical pathways, which in turn regulate the temporal clustering of hippocampal sharp-wave ripples (Clemens et al., 2007; Latchoumane et al., 2017; Staresina et al., 2015).

Accumulating evidence has begun to link the temporal coupling between spindles and slow oscillations during NREM sleep with the overnight stabilization of memories encoded from daytime experiences in humans. This has been demonstrated for procedural (Hahn et al., 2022; Yordanova et al., 2017), spatial (Bastian et al., 2022) and more frequently declarative memory (Denis et al., 2022; Hahn et al., 2020; Helfrich et al., 2018; Mikutta et al., 2019; Muehlroth et al., 2019; Niknazar et al., 2015; Zhang et al., 2020). However, particularly in the context of declarative memory, there is considerable variability in the learning conditions implemented to investigate the relationship with slow oscillation and spindle coupling (Table 1). This has pertinent relevance, as even the behavioural effect of sleep on memory appears to be influenced by the type and strength of learning. While the nature of this relationship has never been systematically tested, evidence exists to suggest that sleep enhances weak associations in memory to a greater extent than strong associations (Diekelmann et al., 2009). However, it has also been proposed that the benefit of sleep for memory follows an inverted-U shaped curve across the depth of encoding (Stickgold, 2009). Similarly, previous studies have demonstrated that pre-sleep learning can influence changes in sleep oscillations. For example, a demanding word-pair learning task has been shown to alter subsequent slow oscillation and spindle activity in central EEG derivations (Mölle et al., 2011, 2009). However, studies have also shown that the nature of the declarative material to be learned, the encoding difficulty, or even inter-individual level in performance, are determining factors in detecting changes in EEG sleep microarchitecture such as sleep spindles after learning (Schabus et al., 2008; Schmidt et al., 2006).

**Table 1.**
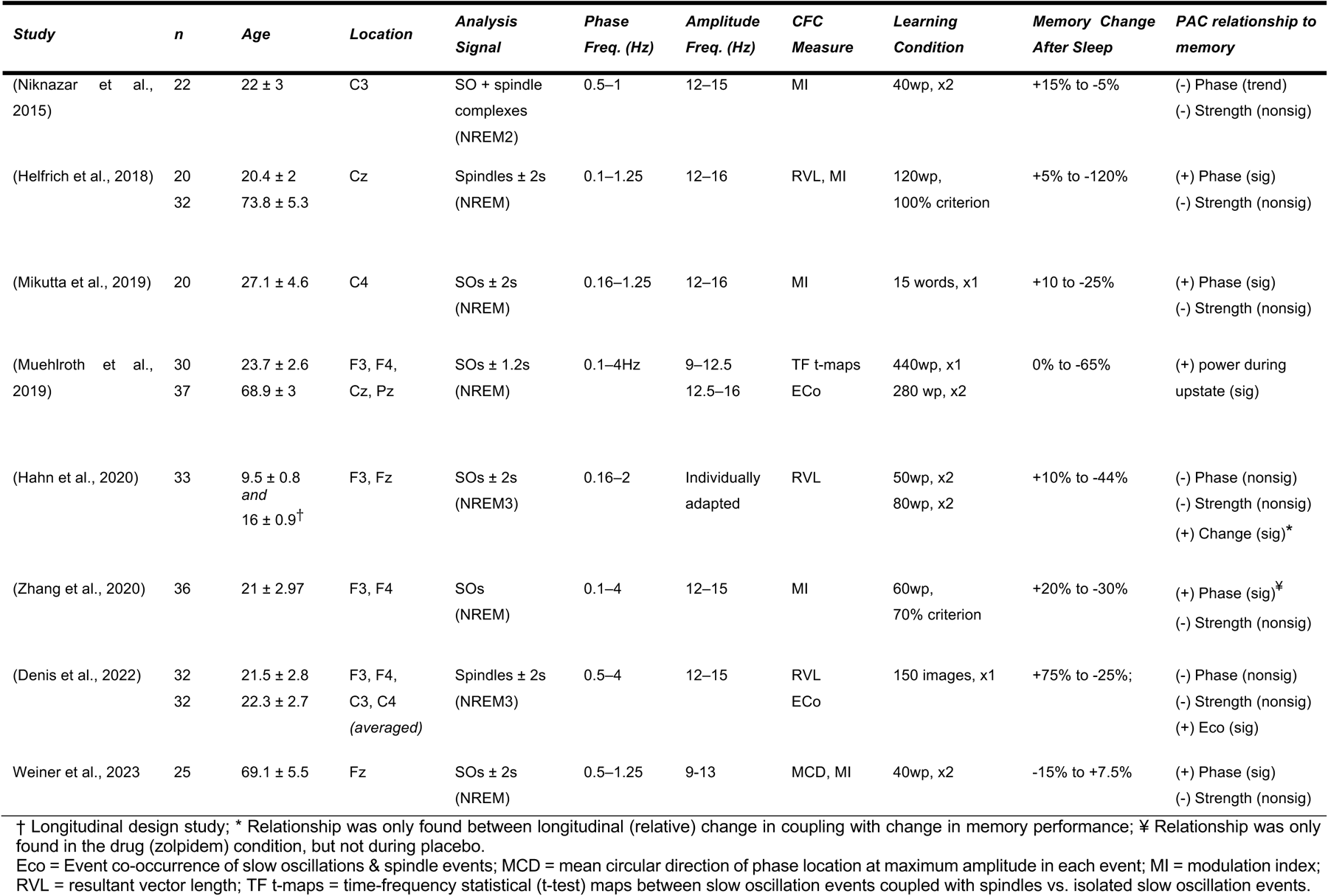
Studies investigating the relationship between declarative memory and phase amplitude coupling of slow oscillations with spindle activity.

Additionally, given their alleged role in reactivation mechanisms for sleep-associated memory consolidation, the extent to which cross-frequency coupling of sleep oscillations fluctuate across nights has received little attention. One study (Cox et al., 2018), found that while there is substantial variation across cortical regions and sleep stages, there is remarkable stability within nights – a characteristic shared by other sleep microarchitecture (Cox et al., 2017; De Gennaro et al., 2005; Finelli et al., 2001; Purcell et al., 2017). It is important to understand whether the coupling between sleep oscillations reflects fluctuating cognitive processes, affected by the amount of active pre-sleep learning efforts and susceptible to interference or enhancement, or whether these are stable individual traits that might expose underlying neurocircuitry. Inter-hemispheric modulation of coupling between slow oscillations and spindles during NREM sleep has been demonstrated in humans following a pre-sleep finger-tapping (motor) task (Yordanova et al., 2017). Yet, the impact of declarative verbal learning prior to sleep on PAC remains unknown. Finally, SO-spindle measures have previously been extracted from variable cortical sites, and the distinction between slow, frontal spindles and fast, parietal spindles in the context of SO-spindle coupling and learning has often been underlooked. As they may be generated from different brain regions and even comprise distinct thalamocortical events (Fernandez and Lüthi, 2020; Marshall et al., 2020), the distinction between these spindle event types should be made when investigating SO-spindle coupling and learning.

This study has two main objectives. Firstly, we measured the strength and timing of PAC between slow oscillation and sigma in the EEG of healthy human participants across two experimental nights (i.e., learning night and control night) to assess intra-individual stability. Specifically, participants were divided into three groups defined by three separate memory tasks with varying load (i.e., the learning paradigm), to test whether the amount or type of verbal word-pair learning prior to sleep influenced subsequent coupling. Secondly, we tested whether the coupling of these rhythms during sleep between the learning period and a delayed recall test predicted overnight changes in performance across the different word-pair learning paradigms. We hypothesised that the type of learning paradigm would affect the relationship between slow oscillation-spindle coupling and overnight changes in memory performance. Given the existence of slow-frontal and fast-parietal sleep spindles (Fernandez and Lüthi, 2020), an exploratory objective was to compare all of the effects on both frontal and parietal electrodes.

## Materials and Methods

### Participants

All participants spoke French or English as a first language. They were recruited through an advertisement in the “students and part-time work” section of an online classified service. Respondents to the advertisement were first taken through a short telephone screening, followed by an in-person semi-structured interview and questionnaires. The telephone and in-person screening ruled out acute and chronic medical conditions, including sleep disorders (e.g., chronic insomnia [sleep difficulty > 3 nights per week for > 3 months], sleep apnea [AHI ≥ 10/h]) and psychiatric disorders (e.g. depression); current use of psychotropic medication or recreational drugs; excessive alcohol or tobacco use; recent (<2 months) travel further than one time zone; current or recent (<1 year) night shift work; and pregnancy. The online screening questionnaires included the Centre for Epidemiological Studies – Depression screening (CES-D)(Radloff, 1977) for depression screening (cut-off at CES-D > 15) and the morningness-eveningness questionnaire (MEQ)(Horne and Ostberg, 1976) to rule out excessively early (MEQ > 70) or late (MEQ < 30) sleep schedules. Out of 259 responders to the online advertisement, 158 were successfully screened by telephone. Among those deemed eligible, 47 were interviewed in person and 38 were screened by overnight PSG to rule out the existence of sleep disorders. Four participants dropped out for personal reasons, 3 for not keeping appointments or for breach of protocol, and 1 because of an allergic reaction to the PSG equipment. Eligible participants (N=41) completed a battery of additional sleep and psychological questionnaires to confirm their eligibility. Sleep disturbances were measured using the Pittsburgh sleep quality index (PSQI, <5) (Buysse et al., 1989) the Epworth sleepiness scale (ESS, <8) (Johns, 1991) and the insomnia severity index (ISI, <8) (Morin et al., 2011). Psychological questionnaires consisted of Beck’s depression inventory (BDI, <10) and the Beck Anxiety Inventory (BAI, <8) (Beck et al., 1996). All participants gave written informed consent prior to the study, which was approved by the Concordia University Human Research Ethics Committee and the Comité d’éthique de la recherche de l’Institut Universitaire de Gériatrie de Montréal (IUGM).

### Experimental design

The experimental design is presented in **Figure 1**. Participants were invited to the sleep laboratory on three non-consecutive nights 1 week apart. The first night served as a polysomnographic (PSG) screening and adaptation night; once cleared of this final screening, participants went on to the experimental phase.

In the experimental phase, participants were divided into three groups that were defined by the word-pair learning paradigms prior to sleep (see *Cognitive Tasks).* Each participant only completed one learning paradigm. Other than the cognitive task performed, the protocols were identical for each group.

The experimental phase consisted of a learning night and a control night, spaced one week apart, and in an order that was counterbalanced between participants. On these nights, participants performed one of two cognitive tasks, a learning task or a non-learning control task, both of them performed on a computer. Tasks were administered two hours before their bedtime such as to provide a 30-minute buffer either side of the participants’ usual bedtime and wake time. They went to bed at their habitual bedtime (no later than 12:30 AM) and slept until they awoke spontaneously (no later than 9 AM). In the morning, participants were offered a light, caffeine-free breakfast 10 minutes after awakening. A final testing (AM) phase was then performed 30 minutes after awakening. Participants were asked to keep regular sleep-wake schedules prior to the study (verified with actigraphy) and to refrain from consuming caffeinated beverages after noon on the recording days. Those who did not maintain regular sleep schedules during the study period were excluded.

### Cognitive tasks

Participants completed the learning task in their first language (English or French). Bilingual participants were asked to choose the language they were most comfortable with. The learning task was a variant of the paired associates task (Plihal and Born, 1999), in which participants were asked to memorize a number of word pair associations. The number of word pairs, and learning trials varied according to the experiment. The first group were presented a list of 40-word pairs. The task finished when participants achieved a correct recall score of ≥60% (24 word pairs), or after five learning trials of the word-pair list (Criterion). The second group were also presented 40 word pairs, however they were only given two learning trials, before the immediate (PM) testing phase was performed (40wp). The third group were presented with 120 word pairs, also shown only twice, before the immediate (PM) testing phase was performed (120wp).

**Figure 1.**
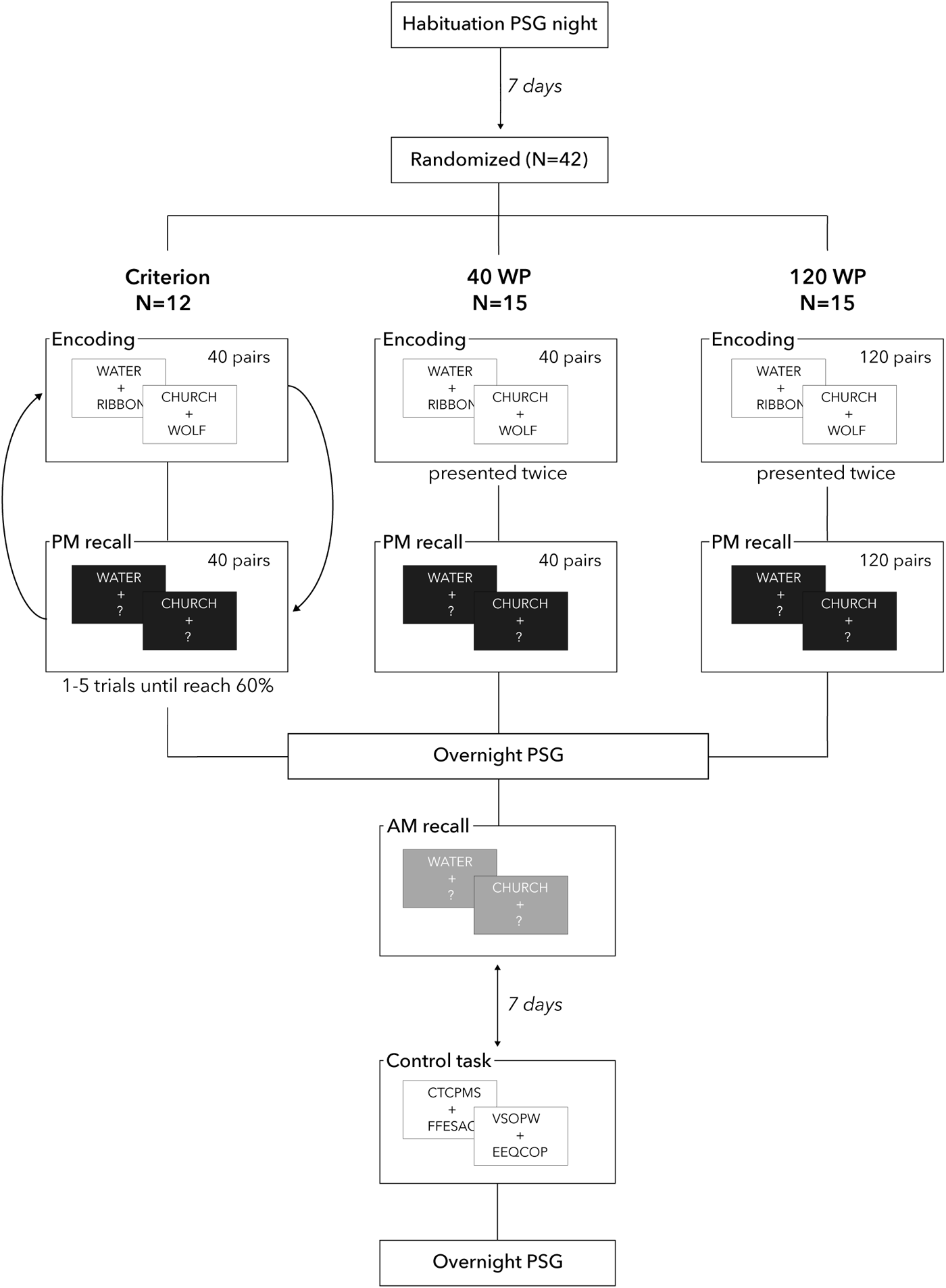
Study design. All participants completed a habituation night in the sleep lab which consisted of a routine polysomnographic (PSG) assessment. Following 7 days, participants completed 1 of 3 experiments, in which the declarative learning paradigm (word pair presentation) was altered. The order of the experimental phase was counterbalanced across participants.

Word pairs were presented sequentially on a computer, and in each learning trail the list was presented in a randomized order. Stimulus presentation for the first trial was 5 seconds, followed by a 5-second blank screen, and a 5-second fixation cross. In all subsequent trials the presentation and breaks were 3 seconds in duration. To control memorization strategies, participants were instructed to use the rest period between each word pair presentation to form a mental image incorporating both words. For instance, if the words were cup and sand, one could imagine a cup filled with sand. All words were French or English nouns of comparable frequency, screened for high concreteness and low emotional salience.

Upon completion of the learning trials, the immediate test phase (PM) was conducted. The first word of each pair was presented onscreen (cue) and participants were asked to verbally recall the associated word (target). Participants had unlimited time to recall the target and received no feedback on their performance. Thirty minutes after awakening the next morning, the test phase was repeated (AM) with the word pair order randomized. Overnight memory performance change was defined as the change in the number of word pairs correctly recalled from pre-to post-sleep, divided by the pre-sleep score.

The control task was based on (Gais et al., 2002) and was designed to resemble the learning task in its duration, structure, visual appearance and difficulty, but without the declarative learning component. Participants were presented with pairs of nonsense alphabetical strings (e.g. BSAVAEC), and were asked to count and orally report the number of letters containing curved lines (e.g. B, C and S, but not A, E and V). For each group, the number of nonsense string pairs in the control task matched the number of word pairs in the learning task. Three test blocks were performed in the evening and one in the morning, matching the learning task.

### Sleep recordings

On the screening night, participants underwent a first full polysomnographic (PSG) sleep recording, including measures of breathing, blood oxygen saturation, electrooculogram (EOG), submental and leg electromyogram (EMG), and electrocardiogram (ECG). On experimental nights, the montage comprised 18 EEG channels, EOG, submental EMG and ECG. Signals were recorded at 512 Hz sampling rate with an online bandpass filter between 0.1 and 128 Hz. Sleep stages were scored offline by experienced raters according to standard AASM criteria (American Academy of Sleep Medicine, 2023). Data segments with EEG artefacts and arousals were also visually identified by an experienced rater and excluded from the analyses. All subsequent analyses of EEG oscillations and CFC were performed on frontal (Fz) and parietal (Pz) electrodes to investigate coupling of slow oscillations with slow-frontal and fast-parietal sleep spindles (Fernandez and Lüthi, 2020). Analyses conducted on a central channel (Cz) are provided in the **Supplementary Materials**.

### Spectral analyses

Power spectral density (PSD) was obtained for all concatenated segments of N2 and N3 sleep over the whole night using Welch’s method, with 4-s Hanning sliding windows overlapped by 2s. The resulting PSD values were log-transformed, given that the underlying distribution of time-windowed EEG power is log-normal (Izhikevich et al., 2018). The PSD values were then averaged across the windowed epochs.

For sigma band analyses, including spindle detection and PAC analyses across NREM sleep epochs, we determined the frequency boundaries of the sigma band for each participant, when a spectral periodic peak was present in the log-log plot of their respective PSDs from the N2 segment. This data-driven determination of participant-specific sigma-band boundaries is robust against the inter-individual variability of spindle peak frequencies (Finelli et al., 2001). We used the *specparam* algorithm to distinguish spectral peaks (Haller et al., 2018) from aperiodic background activity. We observed in our data the typical antero-posterior spindle frequency gradient (Cox et al., 2017; Zeitlhofer et al., 1997) with slower sigma peaks on frontal electrodes (Fz) and faster sigma peaks on posterior electrodes (Pz). We used the highest peak in the 9-13Hz range for frontal electrodes and in the 13-16Hz range for posterior electrodes, to centre the sigma frequency band (with a 4Hz bandwidth).

We used a fixed definition of the slow oscillation frequency band across participants (0.5-1.25 Hz). This range was chosen to cover the band used for the slow oscillation detection, and has been shown to be coupled via PAC to sigma activity (Staresina et al., 2015).

### Event detection

Slow oscillation (SO) and sleep spindle events were detected using previously validated automatic algorithms, implemented in the Wonambi open source toolbox (O’Byrne et al., 2018). The algorithms were run on all artefact-free N2 and N3 epochs, concatenated, and resulting events were reviewed by expert raters. The spindle detector used in the main analyses is based on that used by Mölle and colleagues (Mölle et al., 2011), which involves a thresholding of the root-mean-square (RMS) of the individualised sigma-band signal. Specific to our implementation, we derived different RMS thresholds for each sleep cycle and stage separately to adapt the detection to changing background sigma power over the course of the night. We replicated the analyses using another spindle detection algorithm (Ray et al., 2015) as shown in the Supplementary Materials. The SO detector was based on Staresina and colleagues (Staresina et al., 2015), which involves thresholding the individual’s SO-band signal for events within positive-to-negative zero crossings within the duration criteria. Event duration criteria were 0.5-3 s for spindles, and 0.8-2 s for SOs. We retained the published threshold of ≥75% percentile of SO candidate amplitudes (i.e., the top 25% of events for each participant were tagged as SOs) (Staresina et al., 2015).

### Coupling measures

Spindle-slow oscillation coupling was assessed with three complementary measures: event co-occurrence, preferred couping phase, and the modulation index, as detailed in the following subsections.

### Event co-occurrence measures

A pair of SO and spindle complex were flagged as co-occurrent if they overlapped even partially in time on the same channel (Fz or Pz). To quantify the probability of co-occurrence of SO and spindle events, we used the *intersection-union*, a performance evaluation measure used in the context of information retrieval problems in computer science and machine learning (Rezatofighi et al., 2019). In the context of these problems, *precision* is the fraction of relevant instances among all retrieved instances, and *recall* is the fraction of retrieved instances among all relevant instances. We applied a minimum threshold of 10% overlap and defined all events greater than or equal to this threshold as event co-occurrences. In applying this measure to event co-occurrence, we took one event type (e.g., SOs) as the relevant instances or ground truth, and the other event type (e.g., spindles) as the retrieved instances. We repeated this measure for SOs and spindles separately, to achieve separate sets of SO-spindle complexes (i.e., SO+spindle, SO-spindle) and spindle-SO complexes (i.e., spindle+SO, spindle-SO).

### Event-locked phase-amplitude coupling

We examined phase-amplitude coupling (PAC) between SO (0.5-1.25 Hz) and sigma-band (individually adapted to each participant and electrode) activity. We assesed PAC parameters during the occurrence of SO events only. To create the phase-amplitude distribution, we first extracted signal from each detected event (SO) for the whole night, along with 2 seconds of buffer signal on either side of each event to avoid filter edge artefacts. We then filtered each event in the corresponding low-frequency band (0.5-1.25 Hz) and extracted the instantaneous phase time series using the angle of the Hilbert transform. In parallel, we filtered each event in the (adapted) sigma frequency band and extracted the instantaneous amplitude time series from the modulus of the Hilbert transform. Filtering was done using the tensorpac package (Combrisson et al., 2020), which uses an FIR filter ‘filtfilt’ from Python’s scipy library. Before filtering a padding (extension) of length relative to the filter order was applied to the edges of the signal, to control for filter response given the short time duration of the signal. FIR filters provide crucial advantages over IIR filters when phase relationships are the subject of study as they allow for linear phase (Dvorak and Fenton, 2014). Next, we discarded the buffer signal around each event and binned each value in the amplitude time series by the simultaneous value in the phase (SO) time series (18 bins) to obtain the mean amplitude in each phase bin, producing a single phase-amplitude distribution per SO event. In binning mean amplitudes by phase, we divided the low-frequency cycle into 18 phase bins, balancing precision with robustness (Tort et al., 2010). The mean amplitudes were z-scored across phase bins for each SO event independently, to minimise the influence of amplitude differences prior to all subsequent analyses (Aru et al., 2015; Cole and Voytek, 2017; Gerber et al., 2016). For each SO event, we calculated the phase bin with the maximal mean amplitude of sigma activity. The preferred coupling phase (CP) was calculated as the mean circular direction across all NREM SO events. Finally, the average amplitude across all SO events was calculated independently for each phase bin to calculate one “grand average” phase-amplitude distribution.

### Coupling strength

Measures of the PAC *coupling strength* were measured using the modulation index (MI, (Tort et al., 2010)), a metric based on the Kullback-Leibler divergence, a model-free, information theoretic measure of the distance between two distributions. Briefly, MI measures the level of irregularity in the distribution of the mean amplitudes of a high frequency signal binned by the concomitant phase of a low frequency signal. The MI ranges from 0 to 1, where 0 indicates a uniform distribution of amplitudes across all phase bins, and 1 signifies a total amplitude concentration inside a single-phase bin. In informational terms, it is equal to:

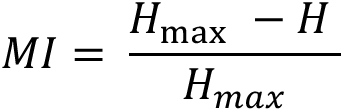

where ***H*** is the Shannon entropy (or randomness) of the phase-amplitude probability distribution and ***H_max_*** is the maximum possible entropy of this distribution (Tort et al., 2010). For all PAC estimations, we followed the recommended guideline that the bandwidth of the frequency for amplitude should be at least double the upper band limit of the frequency for phase (Aru et al., 2015).

We implemented a permutation-based approach to assess the significance of coupling strength against a random phase-amplitude distribution for each subject, as has been implemented previously (Cox et al., 2018). This importantly corrects for the confounding effect of absolute spectral power on absolute coupling strength (MI) that can deviate considerably across electrodes and individuals (Cox et al., 2018, 2017).

Briefly, all the detected SO events for a given channel, night and participant were concatenated into segments of 50 SOs. Any incomplete final segments (i.e., SO count not divisible by 50) were padded with events randomly resampled from the incomplete final segment. For each segment, 400 permutations were performed where the SO phase time series was shuffled with respect to the sigma power time series, and the raw MI was recalculated to create a random distribution of raw MI. The observed MI was then z-scored with respect to this null distribution. The use of segments of identical length for permutation-based z-scoring avoids confounds due to differences in the number of detected SOs. Finally, the z-scores were averaged across all segments to provide the final normalised MI value for each channel, night, and participant. The z-scored measure is a participant- and channel-normalised value, in terms of standard deviations, of the observed coupling strength relative to the average CP estimate under the null hypothesis of no coupling.

We also reran our analyses in two alternative ways to control for the choice of PAC analysis method: 1) calculating PAC only on SO events that were temporally coupled to detected spindle events; and: 2) using the resultant vector length instead of MI (**Supplementary Table 4**).

### Statistical Analyses

Data for the primary outcome measures of interest were screened for univariate outliers, and adherence to statistical assumptions of normality, homogeneity of variance using R. Outliers were inspected using the ‘identify_outliers’ function in R (which detects outliers using boxplot methods). Normality was examined using measures of skewness and kurtosis via a Shapiro-Wilks test. Homogeneity of variance was examined using Levene’s test for each repeated-measure variable pair. Multiple comparisons were corrected using the false discovery rate (FDR) method.

Descriptive statistics were calculated on data pooled across all participants from the 3 learning paradigm conditions. For coupling measures of interest, including the SO-spindle co-occurrence, MI and CP, differences between learning and control nights (‘night effect’) were assessed via a mixed-effects repeated-measures ANOVA with the learning paradigm (criterion, 40wp or 120wp) as the between factor. Post-hoc night effects in CP with circular distributions (i.e., around 0/360°) were calculated with a Watson-Williams test for equal means in a circular distribution, using a permutation shuffling method (10,000 permutations) to facilitate within-subjects comparisons (Watson and Williams, 1956). Watson-Williams tests were only conducted on circular data where a preliminary Rayleigh test (Fisher, 1995) verified the presence of a preferred mean coupling direction (vs. uniform distribution) on both nights. Post-hoc night effects in MI were calculated using linear models. For learning paradigm 1 (criterion), we employed within-subject ANCOVA models to compare MI between learning and control conditions, with the number of trials required to meet the learning criterion of 60% (i.e., between 1-5 trials) as a covariate. For learning paradigm 2 (40wp) and paradigm 3 (120wp), we used paired-samples parametric (T-tests) and non-parametric (Wilcoxon) tests where appropriate. Because the limited sample sizes in each experiment may have concealed any potential significant effects, we also assessed effect size of differences between the Learning and Control nights with Hedge’s ‘g’ to investigate the strength of potential changes. To further quantify the stability of MI and coupling phase between control and learning nights, we used either parametric or nonparametric (linear) correlations or circular-circular correlations and assessed the strength of coefficients.

Finally, we assessed whether the CP or MI were predictive of overnight memory performance change. First, the presence of overnight memory consolidation (i.e., improvement) was confirmed. To determine associations between PAC measures and memory, we used multiple linear regression with overnight memory performance change (%) as the dependent variable and with CP and MI on the learning night as the independent variables. Where a circular distribution of CP across subjects was detected, we first applied a linear transformation to CP before entry into the regression models. The absolute circular distance of each participant’s raw coupling phase (on a scale of 0-360°) to a common reference point (down-state, ‘90°’, for Fz, up-state, ‘270°’, for Pz) was calculated. These resulting values were re-scaled by ±360 where necessary to confine all scores to a range from -180° to +180°. This linear variable reflects a distance either before or after the reference point, and is similar to the one we used in our prior study (Weiner et al. 2023).

## Results

### Participant characteristics

Forty-one healthy young participants aged between 18 and 33 years old were included in this study, including 11 participants who completed the learning paradigm Criterion (22.1 ± 2.9 years old; 6 females), 15 completed the learning paradigm of 40 WP (40 wp, 8 females, 24.9 ± 3.9 years old) and 15 completed the learning paradigm of 120 WP (6 females, 23.6 ± 3.5 years old). There were no significant age differences between learning paradigms. Education ranged from 12 to 22 years. All participants scored within normal ranges in sleep and mood questionnaires (**Supplementary Table 1**). Sleep quality, sleep stage durations, and characteristics of automatically detected spindles and SOs were within normal ranges and did not significantly differ between learning and control nights (**Supplementary Table 2 and 3**).

### Power spectral density

Individualised sigma peak frequencies ranged from 9.1-12.9Hz in the frontal region (Fz channel) and 12.7-14.7Hz in the parietal region (Pz). Peak frequencies did not significantly differ between learning and control nights for any learning paradigm on Fz or Pz (0.5<t<1.8, all p>.05, **Supplementary Figure 1**). Band-limited power across 0.5-30Hz showed no main change between nights for any learning paradigm (all p>.05; **Supplementary Figure 1**). There was no difference between the learning and control night in the offset (0.1<t<1.7, all p>.05) or slope (0.2<t<2.6, all p>.05) of the 1/f exponent) for any learning paradigm, rather these were highly correlated between nights (all r>0.5, p<0.001, **Supplementary Figure 1**).

### Event co-occurrence

Individual and mean percentages of SO events that overlapped in time with spindles during the control night are presented in **Figure 2A**. On average across all three learning paradigms, 12±3% of frontal SO events co-occurred with a slow spindle, while the average number of parietal SOs that co-occurred with fast spindles was similar (11±3%). The percentages were highly correlated between nights (r = 0.72-0.79, p<0.001, **Figure 2B**). There were no night effects in event co-occurrence for either Fz (F = 3.8, p = 0.06, p_FDR_ = 0.240, g = 0.01) or Pz (F=0.26, p = 0.615, g<0.01, **Figure 2C**). A greater proportion of spindles co-occurred with SO events, and this was higher in frontal regions (Fz 32±9%) than in parietal regions (Pz 22±8%, **Figure 2D**). These percentages were also highly correlated between nights (r = 0.73-0.82, p<0.001, **Figure 2E**), and there were no night differences in spindle+SO event co-occurrence on either Fz (F = 1.7, p = 0.197, g = 0.01) or Pz (F = 1.0, p = 0.751, g < 0.01, **Figure 2F**). Post-hoc results for each learning paradigm separately can be found in the **Supplementary Materials**.

**Figure 2.**
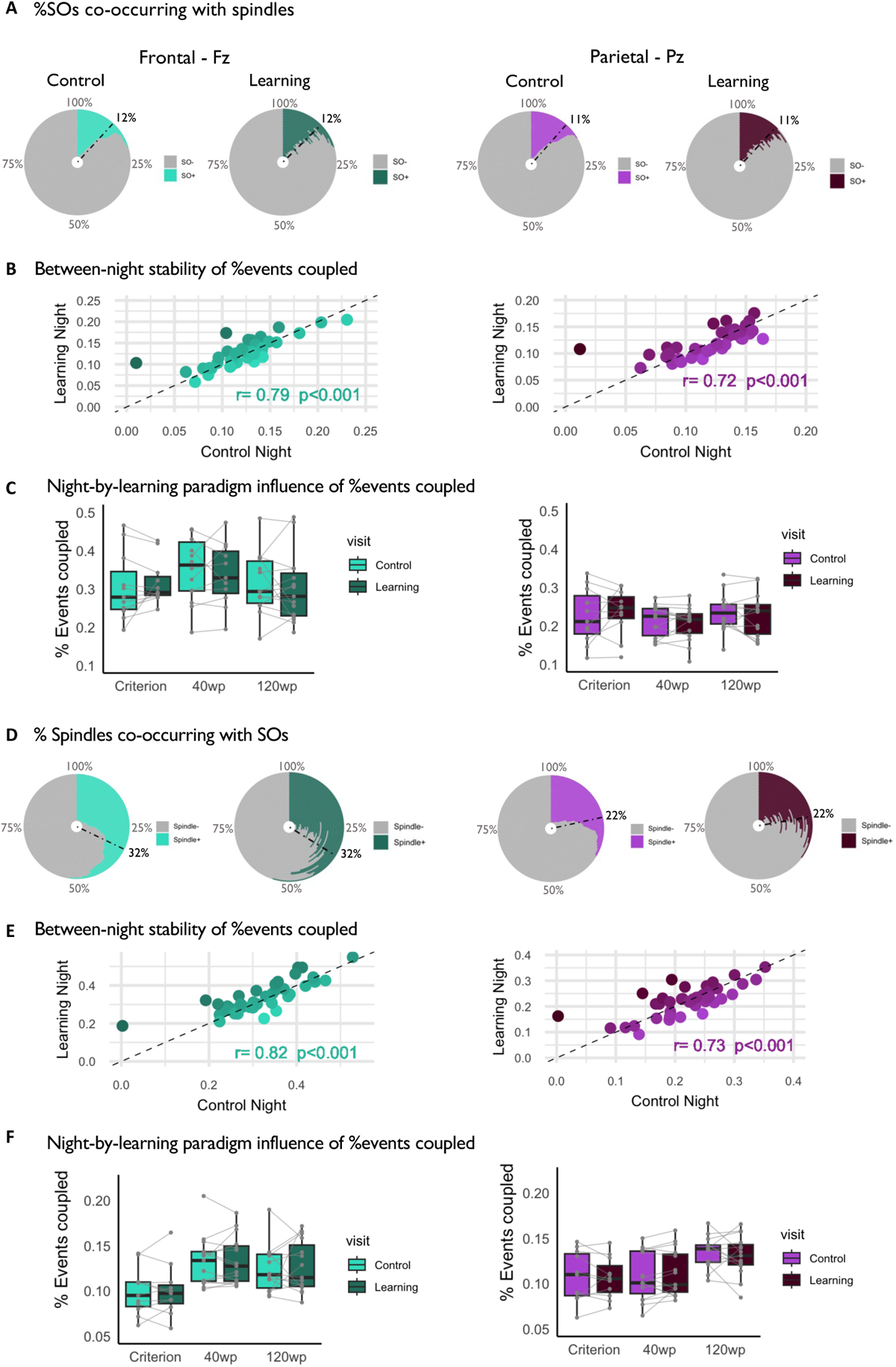
Measures of temporal co-occurrence of slow oscillations (SO) and spindles during NREM sleep. **A**. Race-track plots of individual percentages of SO+spindle complexes on the control across all participants. Each ring represents one participant, ordered by the percentage of coupled events on the control night. Colored sections of rings highlight the percentage of coupled event types; gray sections of rings indicate the percentage of uncoupled events. Black bars represent the average across all participants. **B**. The SO+spindle co-occurrence was highly correlated between nights across all participants. Light colors represent a greater number on the control night, darker colors represent a greater number on the learning night. **C.** The percentage of SO+spindle complexes did not significantly differ between nights for any of the learning paradigms. **D**. Race-track plots of individual and mean percentages of Spindle+SO complexes on the control night. **E.** Between-night correlations of Spindle+SO co-occurrence. Light colors represent a greater number on the control night, darker colors represent a greater number on the learning night. **F.** Percentage of Spindle+SO complexes per night across the 3 experiments. In all plots, results from the frontal region (Fz) are displayed in green and parietal region (Pz) in purple. Light colors represent control night, dark colors represent learning night.

### Effect of pre-sleep learning on phase-amplitude coupling of sleep oscillations

The distribution of sigma power amplitude across the phase of the SO events (averaged over all events and all participants) is displayed in **Figure 3A**. There were no night differences in the mean amplitudes within any phase bin in frontal (Fz) or parietal (Pz) regions. There were also no night differences in the modulation index in Fz (F = 0.36, all p = 0.551, g<0.01). For Pz, there was a small night effect (F = 6.24, p = 0.017, g = 0.02), but this did not pass corrections for multiple comparisons (p_FDR_ = 0.136). Rather, across all participants, the modulation index was strongly correlated between nights for both Fz (r=0.80, p<0.001) and Pz (r=0.75, p<0.001; **Figure 3B**).

**Figure 3.**
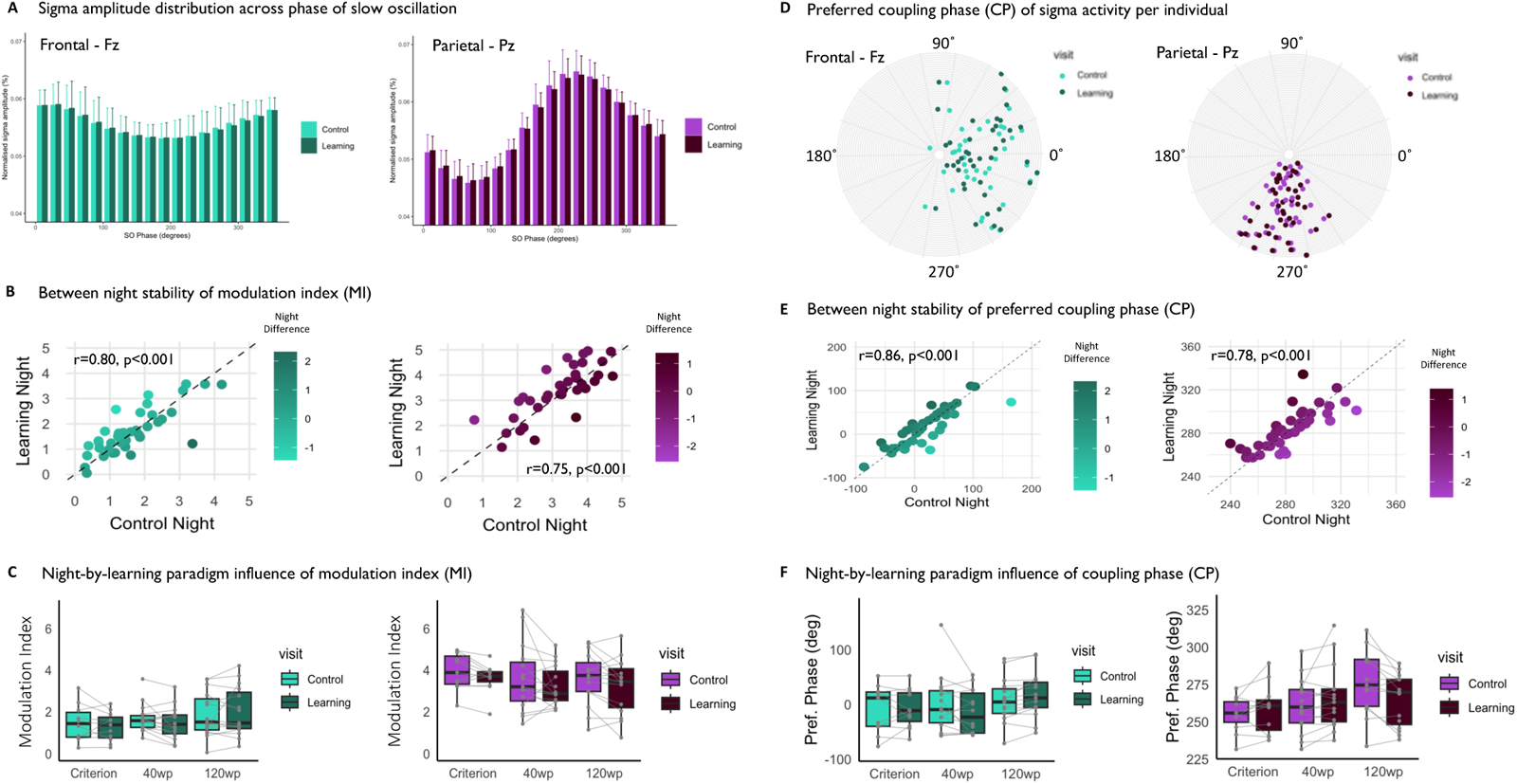
Measures of phase-amplitude coupling between slow oscillations (SO) and spindles during NREM sleep. **A**. Normalised amplitude of sigma frequencies distributed across 18 phase bins of the SO event for all participants. There were no night differences in amplitude within any phase bin. **B**. There were no night differences in normalised strength (modulation index) of SO-spindle coupling per night across the 3 experiments for either frontal (green) and parietal (purple) regions. Between-night correlations of SO-spindle coupling strength measured by the modulation index. Light colors represent a greater number on the control night, darker colors represent a greater number on the learning night. **C**. Circular plots showing the preferred phase of the maximal sigma amplitude per night. Each ring (distance from the centre) represents one participant. **D**. Preferred phase (degrees) of SO-spindle coupling per night across the 3 experiments for frontal (green) and parietal (purple) regions. In all plots: Green = Fz, Purple = Pz; lighter colors = Control night, darker colors = Learning night.

Participant-adapted slow-sigma frequencies were significantly coupled to the up-to-down state phase of frontal SOs, while in the parietal region (Pz) the coupling phase (CP) preferentially preceded the upstate of the SO (**Figure 2C**). The variance of CP values was greater in Fz (sd = 39.7°) than Pz (sd = 18.6°, **Figure 2C**). The CP was significantly correlated between nights for both Fz (r_circ_ = 0.86, p = 0.015) and Pz (r = 0.75, p=0.010; **Figure 3C**). On Fz, there were no night differences in CP (AbsCP, F = 0.12, p = 0.731, g<0.01), and the CP did not shift >20° between nights except for in 2 participants. On Pz there were no night differences in CP (F = 0.12, p = 0.736, g<0.01), and the CP shifted >20° between nights for 1 participant only. There were no night differences in MI or CP when PAC was computed using fixed (uniform 11-16Hz) sigma band frequencies (**Supplementary** Figure 2).

### Phase-amplitude interrelation

There was a significant association between the MI and CP during the control night in both frontal and parietal channels across all participants (**Figure 4A**). This correlation was also observable on the learning night for Pz (**Supplementary Figure 3**). On Fz the relationship was positive (r=0.38, p=0.012), while a negative association was found in Pz (r= -0.64, p<0.001). Specifically, the normalised MI was smaller when the preferred CP was closer to the upstate peak of the SO event. To investigate further what might drive this association, we computed time-frequency spectrograms around the SO events to highlight phase-frequency power patterns. Overall, similar patterns were observed for both Fz and Pz (**Figure 4B**). There was a significant increase in θ, α, and slow-σ power around the SO downstate, while fast-σ and β significantly increased in the down-to-upstate transition and around the upstate of the SO events.

**Figure 4.**
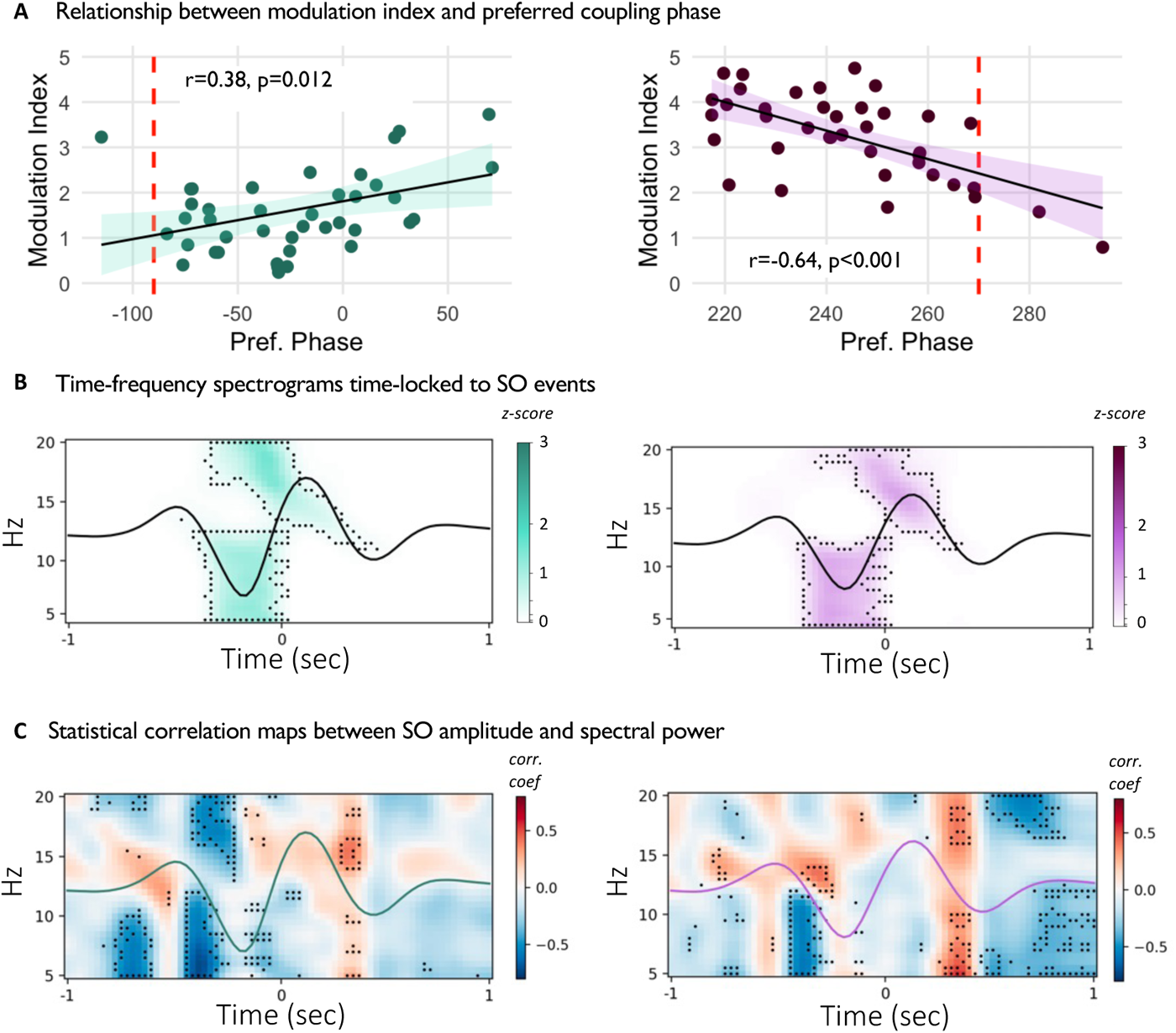
Phase dependent features of coupling strength. **A.** The preferred coupling phase (CP) for each participant was correlated with the magnitude of their modulation index (MI). Specifically, the normalised MI was lower when the CP was closer to the upstate of the slow oscillation (indicated by red-dashed line). Negative phase indicates relative distance to the zero-crossing (ie. 0/360°). **B.** Time-frequency spectrograms time-locked to SO events averaged across SO events and participants (average SO waveform superimposed in black). Activity (power change relative to baseline; see Methods) in the θ, α, slow-σ, and fast-σ ranges is significantly modulated by SOs on both channels. Black outlines represent significant clusters (FDR corrected p<0.05). **C.** Statistical maps illustrating the correlation across participants between the amplitude of the average SO waveform at each timepoint with the average power in the same timepoint across the frequency range (5-20Hz). As a reference, the cohort-average waveform is superimposed. In all plots, Fz is in green, Pz is in purple.

Next, because the MI was smaller when the CP was closer to the upstate, we checked for any relationship between the EEG signal amplitude and any dynamical changes in frequency power across the SO event. Specifically, at every timepoint we correlated the amplitude of the grand-average SO event with the average power in corresponding time-frequency bins across subjects (see Methods). We observed a cluster at the commencement of the SO, such that a steeper decrease to the SO downstate was associated with greater power in in θ, α, and slow-σ frequencies (**Figure 4C**). Significant relationships were also found illustrating that a greater amplitude of the SO upstate was succeeded by more power in fast-σ in Fz, and more power in θ, α and β, but not σ, in Pz. There was no relationship between the SO peak-to-peak amplitude and preferred CP across subjects in either Fz (r= -0.03, p=0.829) or Pz (r = -0.32, p=0.086).

We then assessed the relationship between MI and CP for only the SO events that co-occurred with a sleep spindle (SO+). Firstly, there was no association between CP (across all NREM sleep) and the number of SO+ events on either Fz or Pz. However, SO+ events had a significantly greater MI than SO events that occurred in isolation (SO-), without any differences observed in CP between the two types of SOs (**Supplementary Figure 4A**). On Fz, the significant relationship between MI and CP was only evident for SO+ events (r = 0.4, p=0.009) and not for SO-events (r=0.04, p=0.992). On Pz the relationship was significant for both SO+ (r= -0.67, p<0.001) and SO-(r= -0.67, p<0.001) (**Supplementary Figure 4B**).

Finally, we explored the impact of using individualised adapted sigma frequencies for PAC measures. There was a significant relationship between the inter-individual sigma peak above the 1/f aperiodic activity and the preferred CP (**Supplementary Figure 5**). In both channels, a faster sigma peak correlated with an earlier CP. When fixed sigma frequencies were used, the significant associations between MI and preferred CP were significant for Pz (r = -0.51, p=0.001) but not Fz (r = -0.11, p=0.511, **Supplementary Figure 6**).

### Memory performance

In the first learning paradigm (Criterion) all participants achieved the criterion of 60% accuracy on the word-pairs task, on or before the final (5^th^) presentation of the word list. Performance scores at PM ranged between 60-75%. A sleep-dependent memory consolidation effect was demonstrated by a significant overnight improvement on performance (PM=26±1.9 words, AM=31±2.7 words, t=5.2, p<0.001, g=1.7; **Figure 5A**). The inter-individual range in the change in recall performance (i.e., difference in the number of words correctly recalled at AM and PM) as a percentage of PM performance was 3.5 – 32.1%.

In the 40 WP paradigm, performance scores at PM ranged between 82.5-100%. No sleep-dependent memory consolidation effect was observed (PM=37±2.4 words, AM=37±2.7 words, t=1.0 p=0.33 g=0.09; **Figure 5A**). The range in the improvement in recall performance (i.e., difference between recall at AM and PM) as a percentage of PM performance was -5.5 to +3.0%.

In the 120 WP paradigm, performance scores at PM ranged between 7.5-95.8%. No sleep-dependent memory consolidation effect was observed, conversely there was a trend towards an overnight decrease in memory recall performance (PM=66±32.9 words, AM=65±33.7 words, t=-1.9, p=0.076, g=0.03; **Figure 5A**). The inter-individual range in the improvement in recall performance (i.e., difference in the number of words correctly recalled at AM and PM) as a percentage of PM performance was -15.8 to +11.1%.

**Figure 5.**
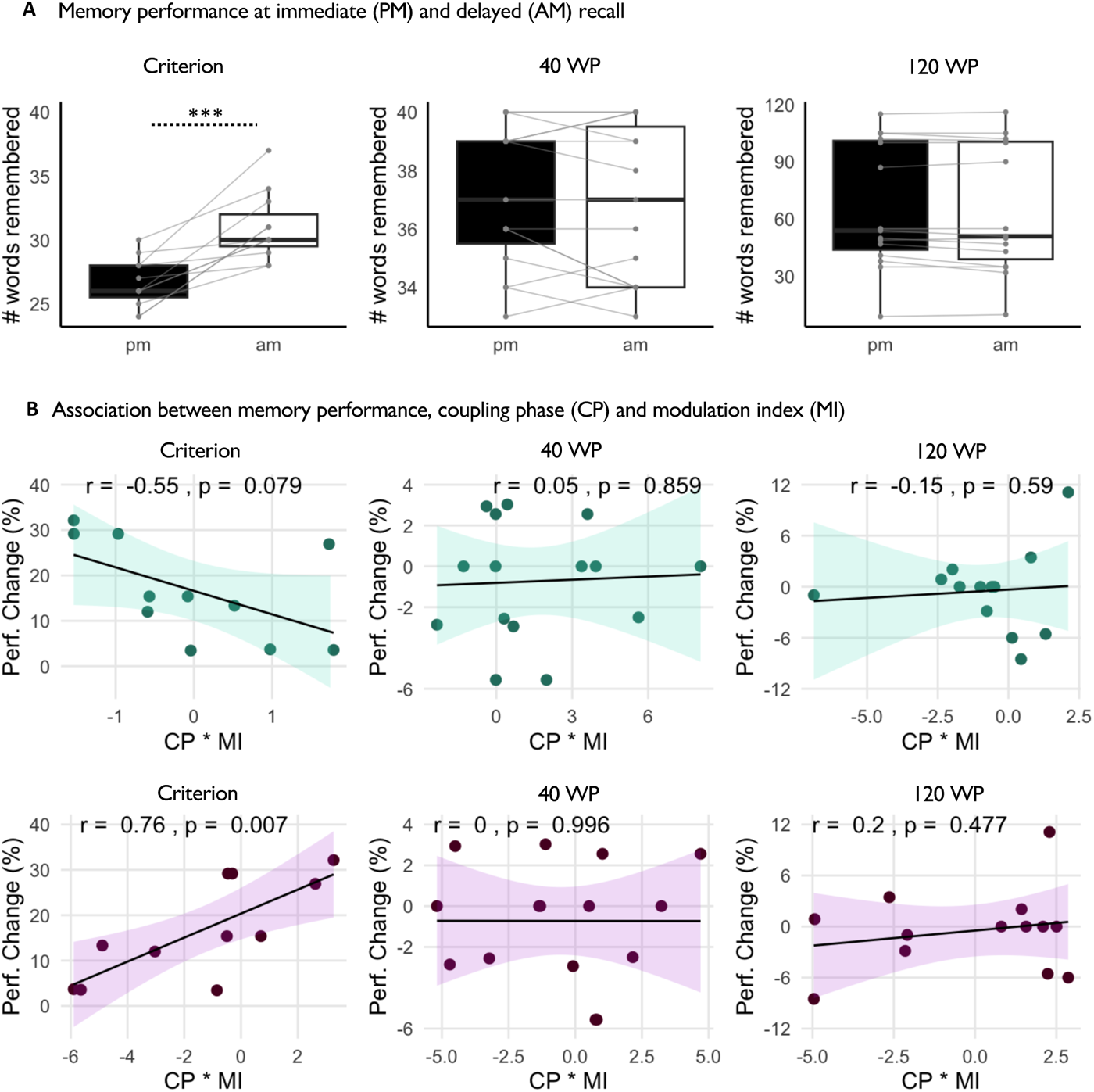
Memory performances for each experiment, where participants had to learn 40 word pairs to a criterion of 60% accuracy before sleep, where presented with 40 word pairs twice, or 120 word pairs twice. **A**. Word recall scores at PM (black) and AM (white) sessions. **B**. Scatterplots of the individual relationships of coupling phase (CP) and modulation index (MI) with overnight memory consolidation for Fz (green) and Pz (purple).

### Memory and phase-amplitude coupling

The significant collinearity between MI and CP invalidated constructing multiple regression models for predicting overnight changes memory with both predictors simultaneously. Instead, we computed a single interaction term consisting of the multiplication of the normalised values of MI and CP (MI*CP), and for any significant prediction we further investigated the contribution of CP or MI with separate exploratory correlations (**Supplementary Figure 7**). For the first learning paradigm (Criterion) there was no significant association between MI*CP and overnight change in memory performance on Fz (r = -0.54, p=0.089, p_FDR_ = 0.236), but there was a significant positive relationship on Pz (r = 0.76, p=0.007, p_FDR_ = 0.043). Exploratory correlations indicated that this was driven by the CP (r = 0.73, p=0.011) more than the MI (r = -0.59, p=0.058). There were no significant associations between MI*CP and overnight changes in memory performance for either of the other learning pardigms (all p>0.477, **Figure 5B**).

## Discussion

This study aimed to assess whether pre-sleep declarative learning can influence the strength and timing of phase-amplitude coupling (PAC) of brain oscillations during subsequent NREM sleep. We measured the PAC of slow oscillations and sigma (spindle) frequency activity in the EEG of healthy young participants across 2 nights (i.e., learning and control nights) to compare intra-individual changes with inter-individual differences. Specifically, three separate experiments with varying cognitive load (i.e., the learning paradigm) were employed to test whether the amount of verbal word-pair learning prior to sleep influenced subsequent coupling. We detected high stability in both the strength and timing of slow oscillation and sigma PAC across nights, with no differences between nights and no detectable influence of varying levels of pre-sleep learning.

Rather, there was much greater inter-individual variability than night-to-night variances within individuals. These results are consistent with recent findings suggesting that NREM slow oscillation-spindle coupling is a stable individual trait (Cox et al., 2018). Several other features of SO (Massimini et al., 2004) and spindle (Cox et al., 2017; De Gennaro and Ferrara, 2003; Ujma et al., 2015) activity also show large, yet reproducible individual differences related to underlying variability in anatomy (Piantoni et al., 2013). Thus, the coupling of neural oscillations during sleep as measured with EEG may reflect neurophysiological processes that are substantially dependent on the integrity of the underlying brain networks, rather than brain states that are influenced by the level of cognitive activity preceding sleep. Other findings showing that the PAC of sleep oscillations are negatively impacted by aging (Helfrich et al., 2018; Muehlroth et al., 2019) and are associated with decreased cortical thickness (Helfrich et al., 2018) are consistent with this hypothesis.

Conventional measures of sleep architecture, such as sleep duration, sleep onset latency, or the number of awakenings after sleep onset, have been shown to fluctuate across nights, being especially worse when sleeping in novel environments (Agnew Jr. et al., 1966; Le Bon et al., 2001). The available literature suggests that neural oscillations may show less night-to-night fluctuation, however within-subject variability of gross sleep architecture and brain oscillations has yet to be systematically and statistically compared. Nonetheless, significantly lower slow wave activity in frontal regions (Mayeli et al., 2022), as well as an interhemispheric asymmetry of slow wave activity in brain networks such as the default mode network (Tamaki et al., 2016) have been shown to fluctuate. These results have also extended to higher sigma and beta activities in medial and left prefrontal areas (Mayeli et al., 2022). Furthermore, changes in spindle activity specific to pre-sleep learning have also only been found in localised regions of the frontal cortex (Schmidt et al., 2006). Thus, it is plausible that any experimental manipulations of PAC are likely to only be found only regionally across the scalp and might explain why we did not detect any differences between nights in this study with a limited montage. With high-density EEG becoming more widely used in sleep research, future studies investigating the relationship between pre-sleep learning and PAC of brain oscillations during sleep would benefit from the superior spatial resolution of implementing this technique into the research design.

We also tested whether the coupling of these rhythms was related to overnight changes in performance across different word-pair learning paradigms, to assess their relationship with different processes of memory consolidation or stability. We found that there was a significant relationship between PAC of slow-oscillations and spindles, only in the learning paradigm that demonstrated an overnight consolidation effect (i.e., participants significantly improved their memory performance from immediate to delayed recall). In the other learning paradigms, neither individual nor group level performances significantly changed between immediate and delayed recall, which could explain why no relationship was observed with PAC of brain oscillations during sleep. While learning refers to the accumulating performance improvements within a given training experience (within-session gains), memory consolidation refers to the acquisition of delayed (between-session) gains in performance that can occur in the absence of any additional practice (Brashers-Krug et al., 1996; Hauptmann et al., 2005). Previous studies have observed significant associations between PAC measures and performance on declarative memory (word pair) tasks without observing strong consolidation effects (**Table 1**). However, as we have recently discussed (Weiner et al., 2023), there are noticeable differences in experimental and analytic methods across these studies that are important to consider when reviewing the currently available data. Relevantly, this was the first time that the relationship between PAC and declarative memory has been directly compared across different learning paradigms using the same analysis method to extract measures of PAC. Alternatively, some significant relationships have been reported with the preferred phase of SO-spindle coupling when there was a decrease in performance at delayed recall (i.e., forgetting), indicating that coupling of brain oscillations during NREM sleep might also be protective of memory decay. While none of the learning paradigms in this study produced any effects of forgetting, we cannot rule out that SO-spindle PAC might be involved in sleep-dependent forgetting. Instead, we were able to demonstrate that PAC of sleep oscillations are specifically related to sleep-dependent memory consolidation compared to pre-sleep learning ability.

These findings reiterate the importance of considering the type of learning paradigm when designing experiments for assessing the role of sleep for memory. This is particularly critical when interpreting the type of memory process in question (e.g., encoding, consolidation, forgetting) that might be associated with sleep and especially brain oscillations during sleep. It has been proposed that the outcome of memories following sleep may depend on their relative strength or stability at sleep onset (Talamini et al., 2008). While one study found that sleep enhanced the associative strength of declarative memories more robustly for weaker associations (Drosopoulos et al., 2007), other studies have shown that sleep has a tendency to enhance stronger associative memories based on recency of encoding (Talamini et al., 2008) or depth of encoding (Schabus et al., 2008). This has led some to propose that the benefits of sleep for memory follow inverted-U shaped curve across the depth of encoding (Stickgold, 2009). A recent study also reported that that age deficits in sleep-dependent memory consolidation were more prominent for items of medium encoding quality (Muehlroth et al., 2020). Nonetheless, these results support the notion of Schabus and colleagues (2008) that memory must have reached sufficient encoding depth during learning for offline processing during sleep to become evident, as we only found a consolidation effect on performance when participants learned to criterion. Finally, more efficient learning ability (high performance prior to sleep) has also been shown to exhibit greater effects of sleep on changes in memory performance (Schabus et al., 2008; Tucker and Fishbein, 2008). Our findings assert the importance of learning paradigm on this effect also, as the results were opposite for learning to criterion vs fixed exposure. While not significant, in the learning to criterion condition, we observed a trend for the bottom 50% of performers at immediate recall to benefit more during the intervening sleep period than the high performers (**Supplementary Figure 8**).

Contrarily, in the other two learning paradigms the trend was in the opposite direction, especially in the condition with 120 word-pairs where the low performers had significantly greater negative gains (i.e., forgot more words) than the high performers. However, there were no associations with SO-spindle PAC based on performance on the immediate recall. Instead, these findings demonstrate that the learning paradigm is more important for revealing relationships between memory consolidation and PAC of brain oscillations during NREM sleep.

Finally, we observed dependencies between the commonly used metrics of SO-spindle PAC. To our knowledge, we are the first to report any associations between these metrics. There were significant associations between the strength (modulation index, MI) of coupling and the preferred coupling phase (CP) across individuals. Generally, the MI was lower when the CP was closer to the upstate. Specifically, in the frontal region a later CP was associated with a greater MI, while in the parietal region individuals with an earlier CP exhibited a greater MI. This is consistent with the known dynamics of SO coupling with slow and fast spindles, as it has been demonstrated that slow frontal spindles are coupled to the up-to-downstate transition of SO events, while fast parietal spindles preferentially couple just preceding the upstate (Cox et al., 2018). This is also consistent with the patterns we observed in the time-frequency spectrograms and the relationship between CP and the individualised sigma spectral peak. In the frontal cortex a slower sigma peak frequency of spindles suggests coupling later in the SO event, closer to the downstate where there is a robust increase in spectral power of frequencies below 10Hz (Figure 4). It is possible that given some individuals in this study had slow sigma peaks below 10Hz, there could be contamination of alpha activity in the PAC analyses. This would explain that when using uniform (higher, 11-16Hz) frequency bands, the relationship between MI and CP was no longer significant in the frontal region (**Supplementary** Figure 6). However, when using adapted bands PAC metrics in frontal regions exhibited strong fingerprint-like effects (i.e., large inter-individual differences & stable within-subject consistency). Additionally, this does not explain why a preferred CP closer to the SO upstate was negatively associated with a reduced MI for fast parietal spindles. This was unlikely influenced by the number of spindles that co-occurred with SO events, as although the MI was greater, there was no difference in CP between SO+ and SO-events. Nevertheless, the dependency between MI and CP is an important consideration for studies wishing to compare the importance of both PAC strength and timing in NREM sleep. Overall, these results indicate that the relationship between preferred coupling phase (timing) and modulation index (strength) of SO-spindle coupling is complex and dependent on multiple factors.

## Conclusion

Here we reported on the phenomenology of PAC between typical NREM oscillations related to various declarative learning paradigms that differed on the amount of information learned and the frequency of exposure to the information. We found that SO-sigma PAC is unaffected by learning regardless of the learning paradigm, but that it is only associated with delayed recall of information when there exists successful memory consolidation. These findings further suggest that PAC is a stable individual brain trait. This is relevant for the continued investigation of PAC in the context of disease or aging, as changes in the coupling of endogenous brain oscillations during sleep may more likely be a marker of degraded network interactions rather than represent fluctuating processes (ie. sleep quality or cognition).

## Acknowledgements

We would like to acknowledge Claire André and Eden Debellemaniere for their assistance with data collection, and Sylvain Baillet for his comments on this manuscript.

## Supplementary Materials

**Supplementary Figure 1.**
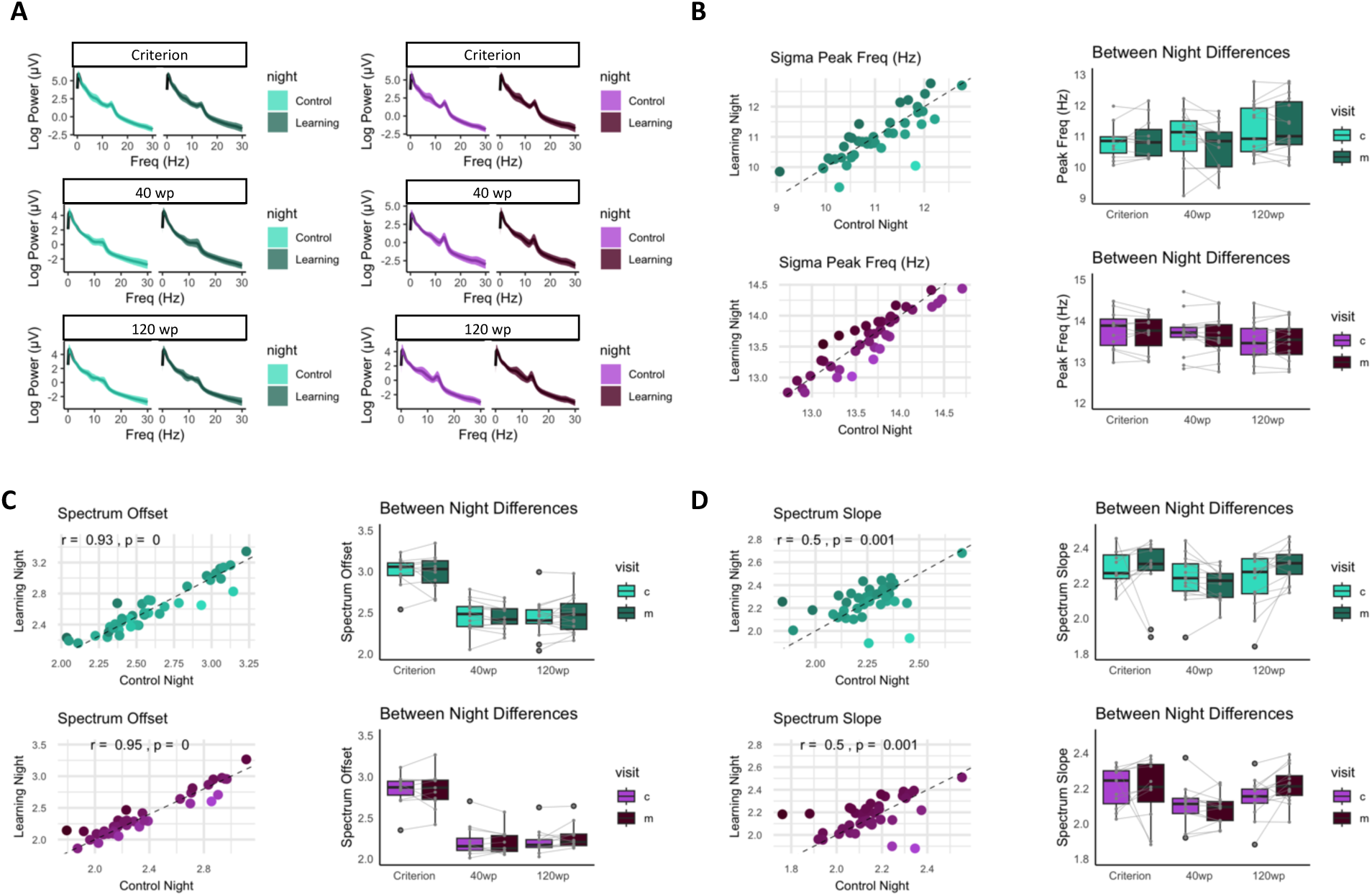
**A** Power spectral activity between control and learning nights for both frontal (Fz, green) and parietal (Pz, purple) regions across the three learning conditions. There were no between-night differences in power for any frequencies across learning conditions. **B** The individual sigma spectral peak (above the aperiodic 1/f activity) for each participant was significantly correlated between nights, and there were no between night differences. **C** The individual spectral offset (intercept of aperiodic 1/f activity) for each participant was significantly correlated between nights, and there were no between night differences. **D** The individual spectral slope (slope of aperiodic 1/f activity) for each participant was significantly correlated between nights, and there were no between night differences.

**Supplementary Figure 2.**
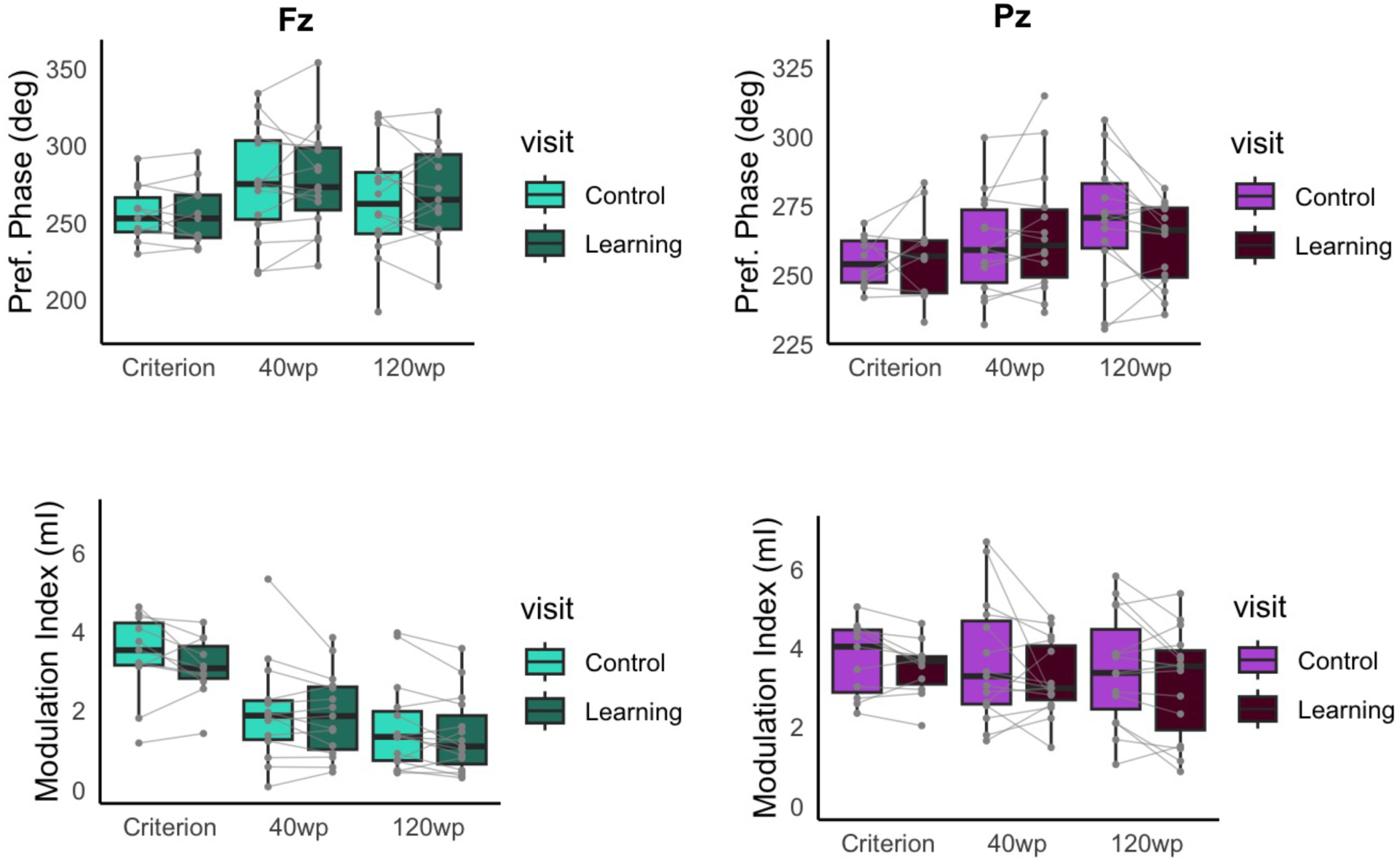
Box plots of preferred phase (top panel) and modulation index (bottom panel) for both frontal (Fz, green) and parietal (Pz, purple) regions across learning conditions when PAC was computed using fixed (uniform 11-16Hz) sigma band frequencies. There were no between night differences in any learning condition.

**Supplementary Figure 3.**
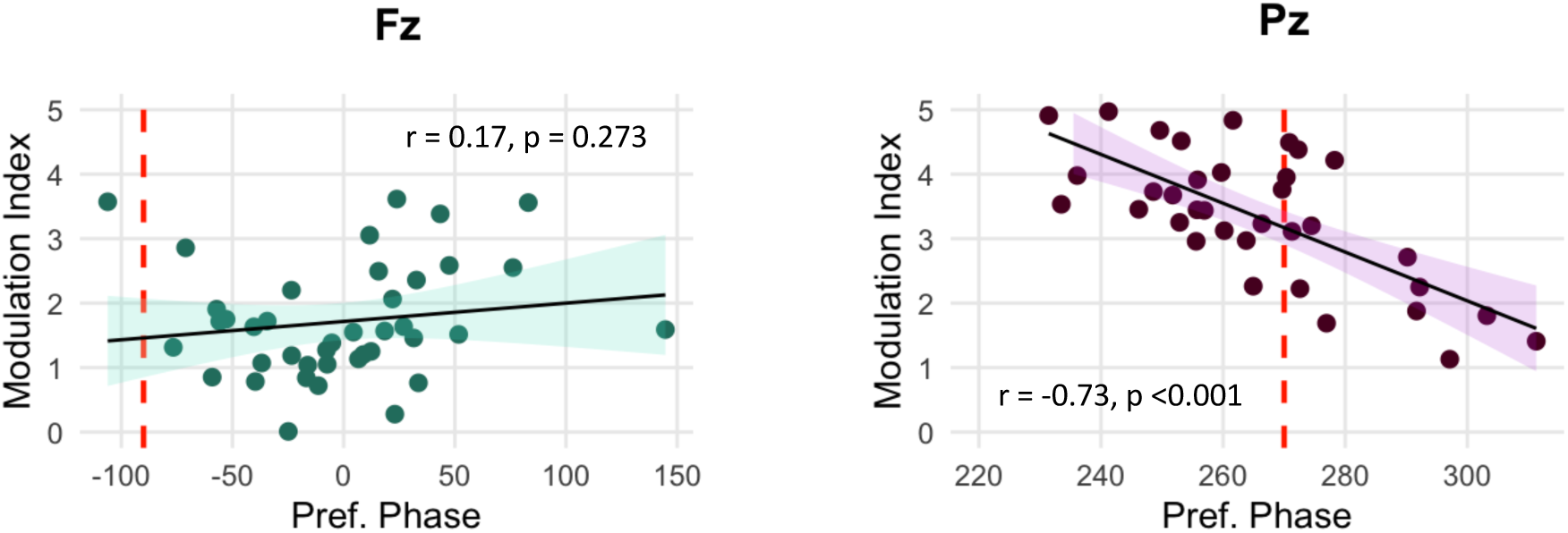
**A** The relationship between the modulation index and preferred coupling phase was only significant for Pz (purple) during the control night.

**Supplementary Figure 4.**
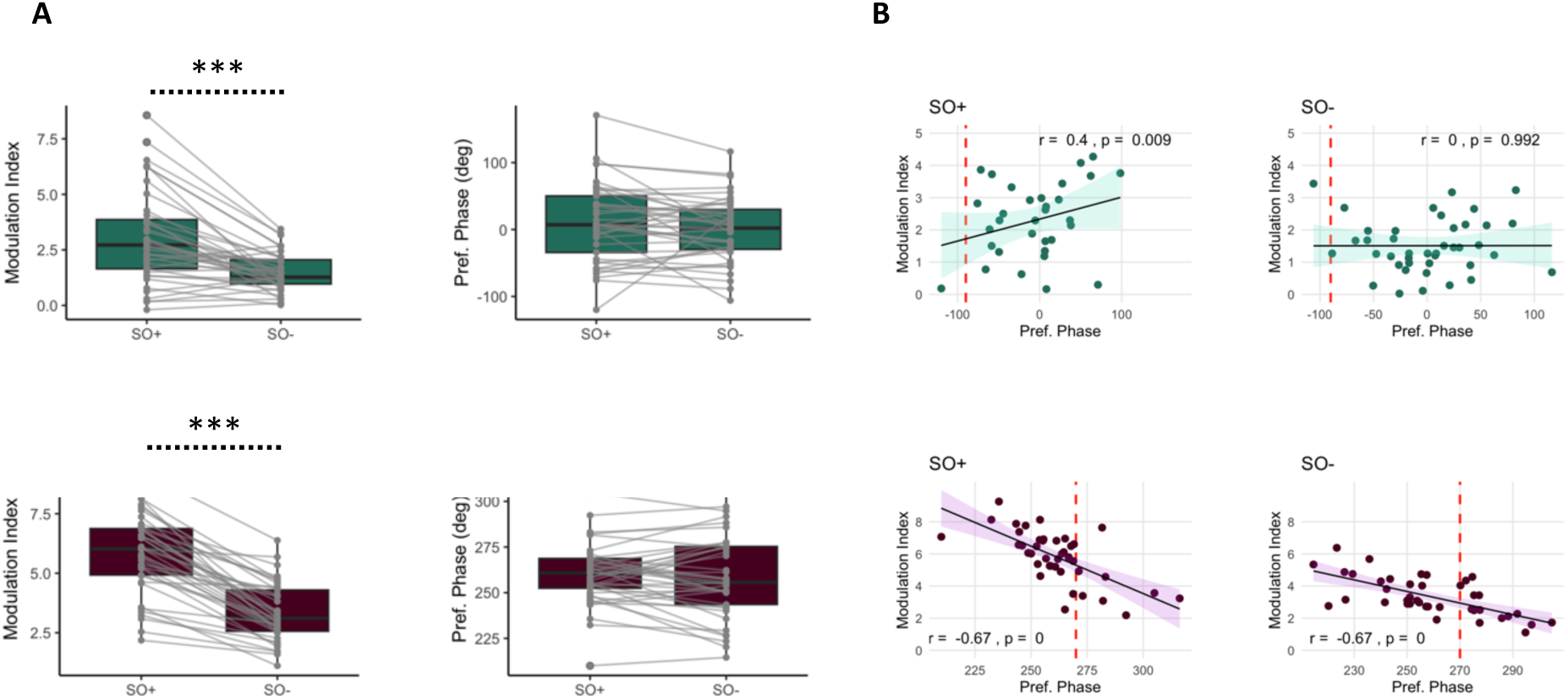
**A** Differences in modulation index (left) and preferred phase (right) for SO events that co-occur with spindles (SO+) or occur in isolation (SO-) for frontal (Fz, green) and parietal (Pz, purple) regions. **B** The relationship between the modulation index and preferred coupling phase was only significant for SO+ and not for SO-in frontal regions, while it was significant for both SO+ and SO-in parietal regions.

**Supplementary Figure 5.**
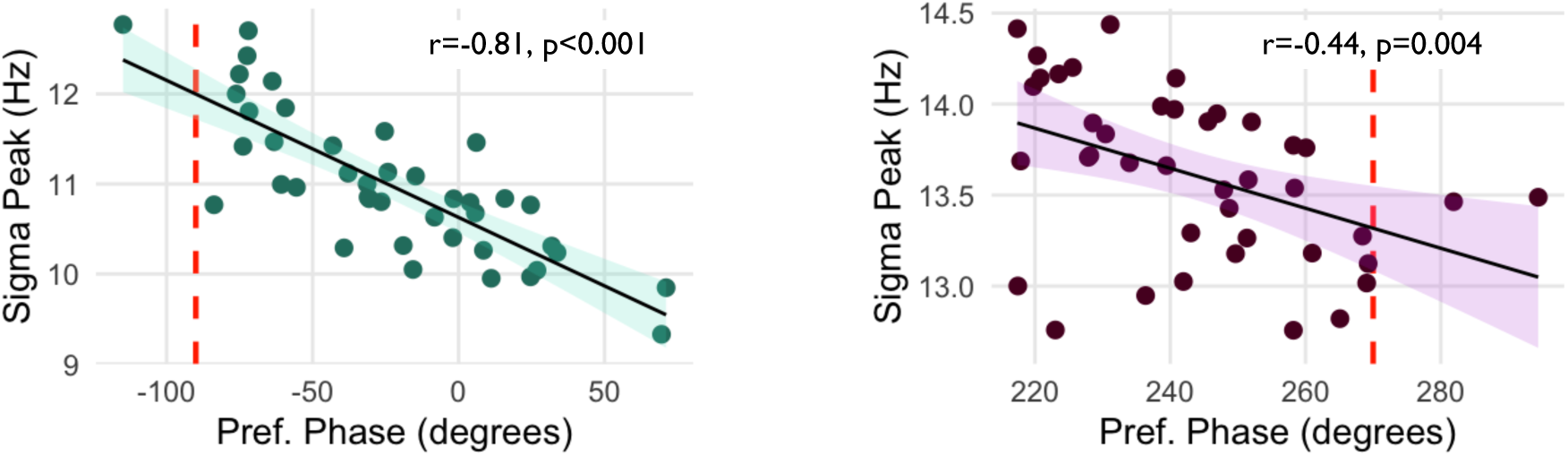
The relationship between the individual sigma spectral peak (above the aperiodic 1/f activity) and the preferred phase of slow oscillation and spindle coupling for frontal (Fz, green) and parietal (Pz, purple) regions. Frontally, the sigma peak frequency was faster when the CP was closer to the upstate of the slow oscillation (indicated by red-dashed line). In parietal regions, the sigma peak frequency was slower when the CP was closer to the upstate of the slow oscillation.

**Supplementary Figure 6.**
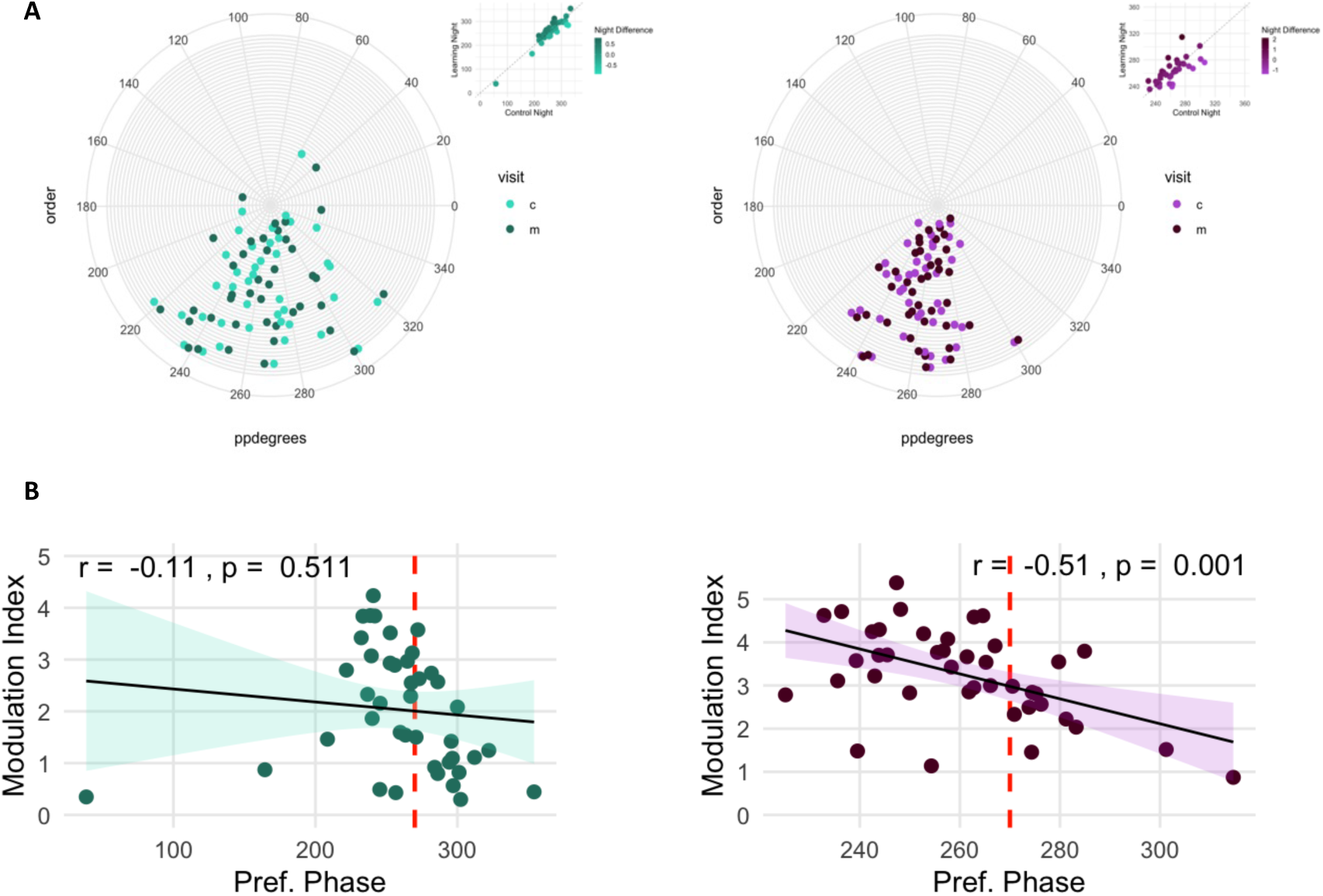
**A** The preferred coupling phase (CP) of slow oscillation spindle coupling when sigma frequency bands (11-16Hz) were uniform across all participants for the phase amplitude coupling analyses for frontal (Fz, green) and parietal (Pz, purple) regions. The CP remained consistent with the analyses using adapted bands for parietal regions, while in frontal regions the CP was more clustered around the upstate (rather than in the up-to-downstate transition with adapted bands). **B** The relationship between CP and the modulation index (coupling strength) was no longer significant in frontal regions when uniform sigma frequency bands (11-16Hz) were implemented but remained significant in parietal regions.

**Supplementary Figure 7.**
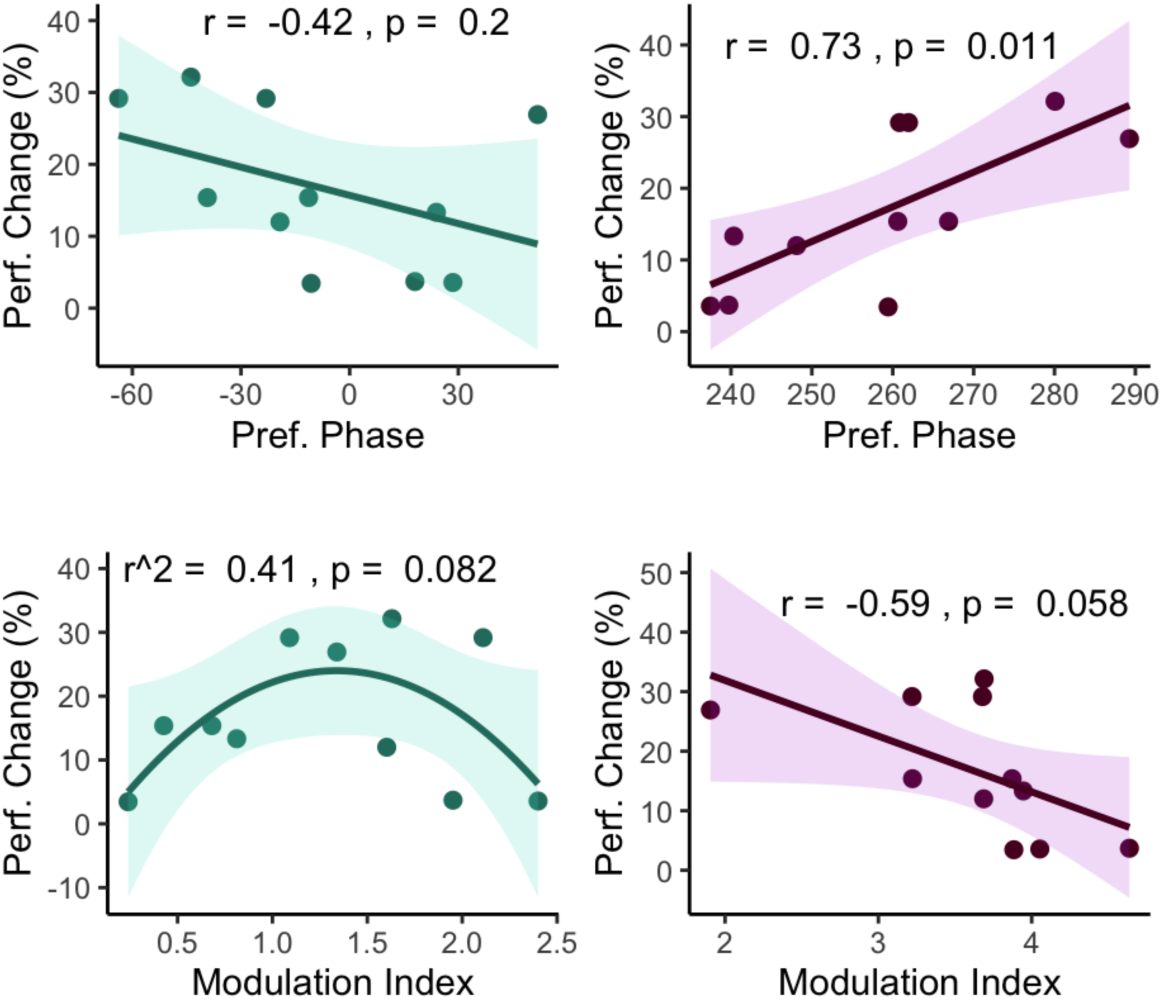
Scatterplots of the associations between preferred coupling phase (top panel) and modulation index (bottom panel) for frontal (Fz, green) and parietal (Pz, purple) regions in the learning to criterion paradigm.

**Supplementary Figure 8.**
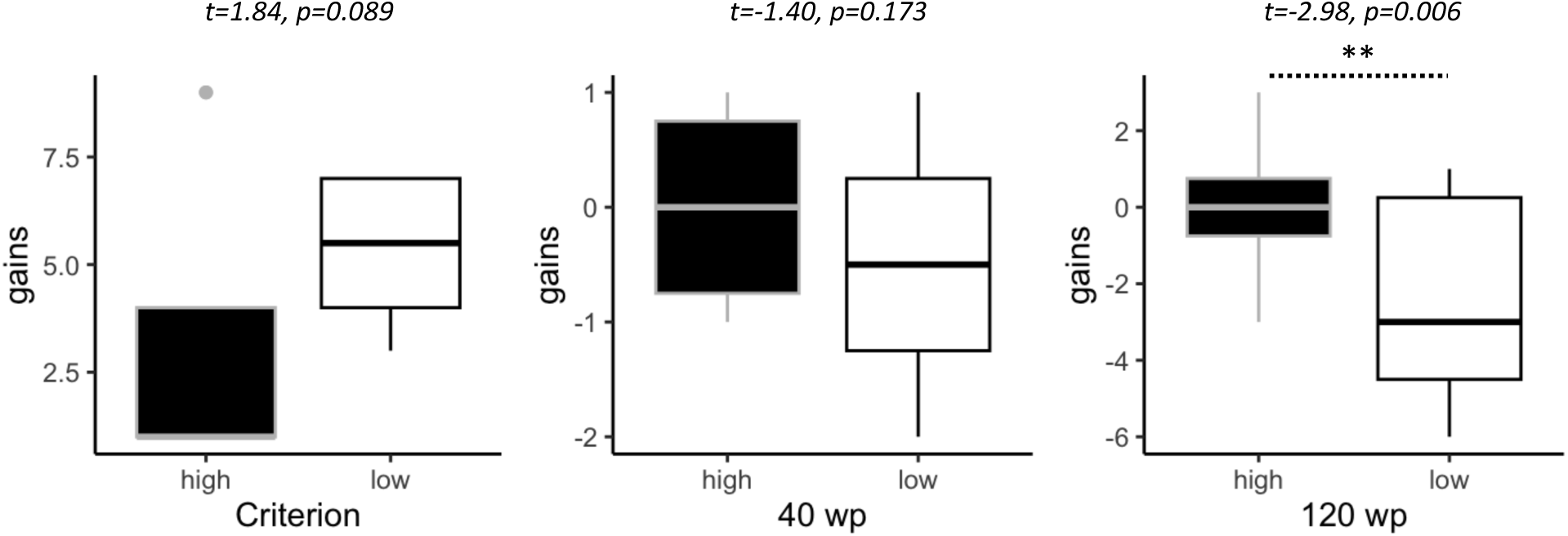
Differences in overnight performance gains (the difference in the number of words recalled between immediate and delayed recall tests) on the word-pair learning task for high and low performers. Groups were based on a median split of performance (the number of words recalled) at immediate recall.

## Supplementary Results

### Event co-occurrence per learning paradigm

For the learning to criterion paradigm, 9.9±2.6% of frontal SO events co-occurred with a slow spindle, while the average number of parietal SOs that co-occurred with fast spindles was similar (10.8±2.8%). The percentages were highly correlated between nights (r = 0.79-0.88, p<0.01). There were no night effects in event co-occurrence for either Fz (t = -0.17, p = 0.868, g = -0.02) or Pz (t = 0.34, p = 0.737, g = 0.05). A greater proportion of spindles co-occurred with SO events, and this was higher in frontal regions (Fz 30.5±8.9%) than in parietal regions (Pz 22.5±7.1%). These percentages were also correlated between nights (r = 0.66-0.65, p<0.05), and there were no night differences in spindle+SO event co-occurrence on either Fz (U = 28, p = 0.689, g = -0.11) or Pz (t = -0.69, p = 0.506, g = -0.16).

For the 40WP paradigm, 13.7±3.0% of frontal SO events co-occurred with a slow spindle, while the average number of parietal SOs that co-occurred with fast spindles was similar (13.7±3.0%). The percentages were highly correlated between nights (r = 0.79-0.89, p<0.001). There were no night effects in event co-occurrence for either Fz (t = -0.25, p = 0.806, g = -0.03) or Pz (t = -0.37, p = 0.717, g = -0.05). A greater proportion of spindles co-occurred with SO events, and this was higher in frontal regions (Fz 35.5±8.0%) than in parietal regions (Pz 20.9±5.1%). These percentages were also highly correlated between nights (r = 0.86-0.91, p<0.001), and there was a night difference in spindle+SO event co-occurrence on Fz (t = -2.4, p = 0.034, g = -0.24), but this did not pass corrections for multiple comparisons (p_FDR_ = 0.136). There were no night differences in spindle+SO event co-occurrence on Pz (t = -0.18, p = 0.862, g = -0.02).

For the 120WP paradigm, 12.2±4.0% of frontal SO events co-occurred with a slow spindle, while the average number of parietal SOs that co-occurred with fast spindles was similar (13.1±4%). The percentages were correlated between nights (r = 0.54-0.67, p<0.05). There were no night effects in event co-occurrence for either Fz (t = -1.47, p = 0.163, g = -0.29) or Pz (t = -0.43, p = 0.677, g = -0.11). A greater proportion of spindles co-occurred with SO events, and this was higher in frontal regions (Fz 29.9.5±1.0%) than in parietal regions (Pz 22.6±8.0%). These percentages were also correlated between nights (r = 0.71-0.73, p<0.01), and there were no night differences in spindle+SO event co-occurrence on either Fz (t = -0.89, p = 0.391, g = -0.16) or Pz (t = 0.08, p = 0.937, g = 0.02).

### Phase-amplitude coupling on central scalp regions (Cz)

Individual and mean percentages of SO events that overlapped in time with spindles are presented in Supplementary Figure 9A. On average across all three learning paradigms, 12±3% of frontal SO events co-occurred with a slow spindle. The percentages were highly correlated between nights (r = 0.81, p<0.001, **Figure 2B**). There were no night effects in event co-occurrence (F = 3.8, p = 0.06, p_FDR_ = 0.240, g = 0.01, **Figure 2C**). A greater proportion of spindles co-occurred with SO events (26±8%, **Figure 2D**), and these percentages were also highly correlated between nights (r = 0.75, p<0.001, **Figure 2E**). Finally, there were no night differences in spindle+SO event co-occurrence (F = 1.7, p = 0.197, g = 0.01, **Figure 2F**).

**Supplementary Figure 9.**
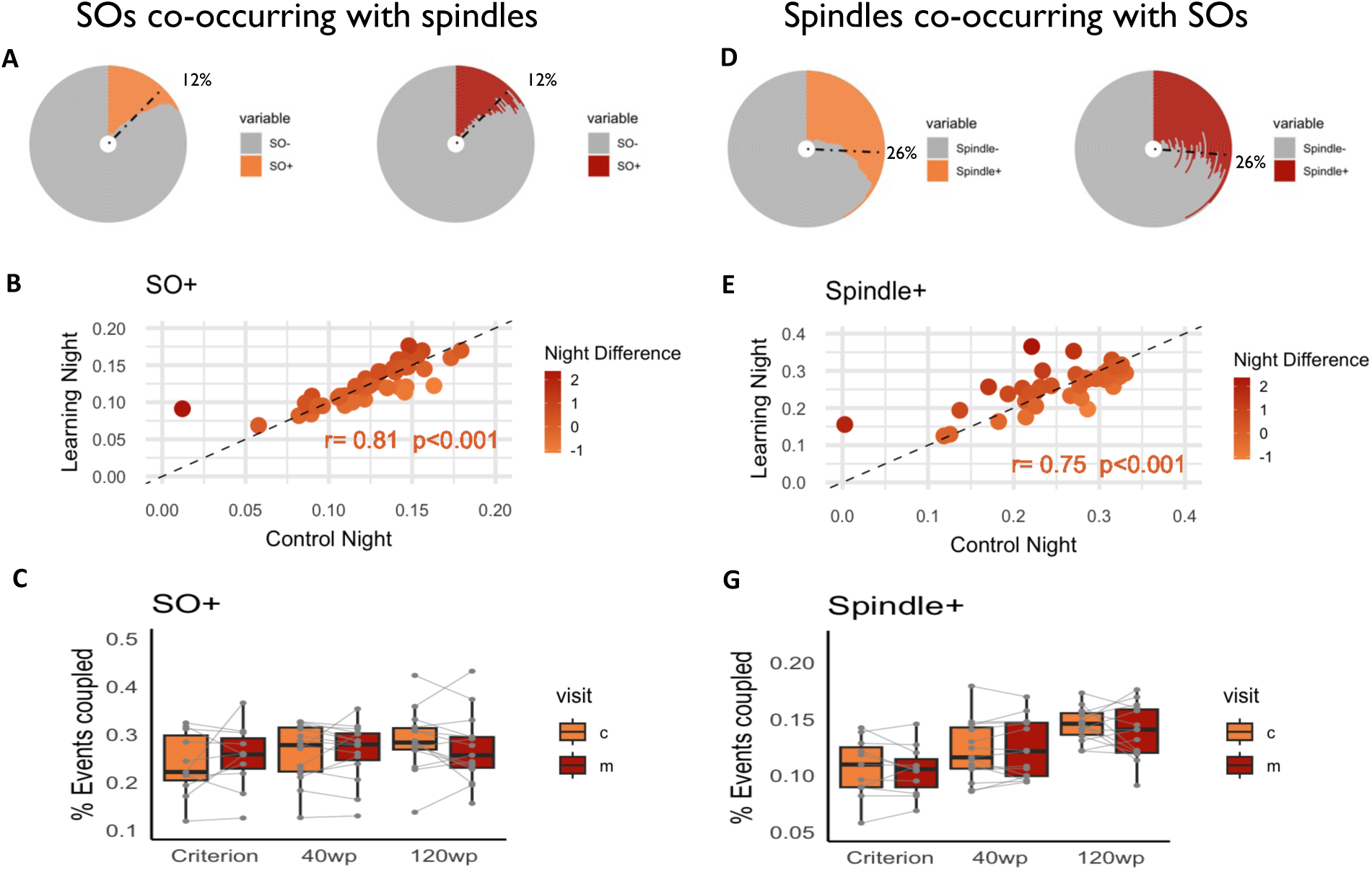
Measures of temporal co-occurrence of slow oscillations (SO) and spindles during NREM sleep on the central recording site (Cz). **A**. Race-track plots of individual percentages of SO+spindle complexes on the control across all participants. Each ring represents one participant, ordered by the percentage of coupled events on the control night. Colored sections of rings highlight the percentage of coupled event types; gray sections of rings indicate the percentage of uncoupled events. Black bars represent the average across all participants. **B**. The SO+spindle co-occurrence was highly correlated between nights across all participants. Light colors represent a greater number on the control night, darker colors represent a greater number on the learning night. **C.** The percentage of SO+spindle complexes did not significantly differ between nights for any of the learning paradigms. **D**. Race-track plots of individual and mean percentages of spindle+SO complexes on the control night. **E.** Between-night correlations of Spindle+SO co-occurrence. Light colors represent a greater number on the control night, darker colors represent a greater number on the learning night. **F.** The percentage of Spindle+SO complexes did not significantly differ between nights for any of the learning paradigms.

The distribution of sigma power amplitude across the phase of the SO events (averaged over all events and all participants) is displayed in **Supplementary Figure 10A**. There were no night differences in the mean amplitudes within any phase bin. There were not any night differences in the modulation index in Cz (F = 0.36, all p = 0.551, g<0.01, **Supplementary Figure 10C**). Rather, across all participants, the modulation index was strongly correlated between nights (r=0.80, p<0.001, **Supplementary Figure 10B**).

Participant-adapted sigma frequencies on Cz were significantly coupled to the upstate of the SO (**Supplementary Figure 10D**). The values and variance of CP were similar to those observed in Pz, indicating that fast spindle coupling was more dominant in central regions, as expected. The CP was significantly correlated between nights (r = 0.77, p<0.001; **Supplementary Figure 10E**). There were no night differences in CP (F = 0.12, p = 0.736, g<0.01), and the CP shifted >20° between nights for 1 participant only.

**Supplementary Figure 10.**
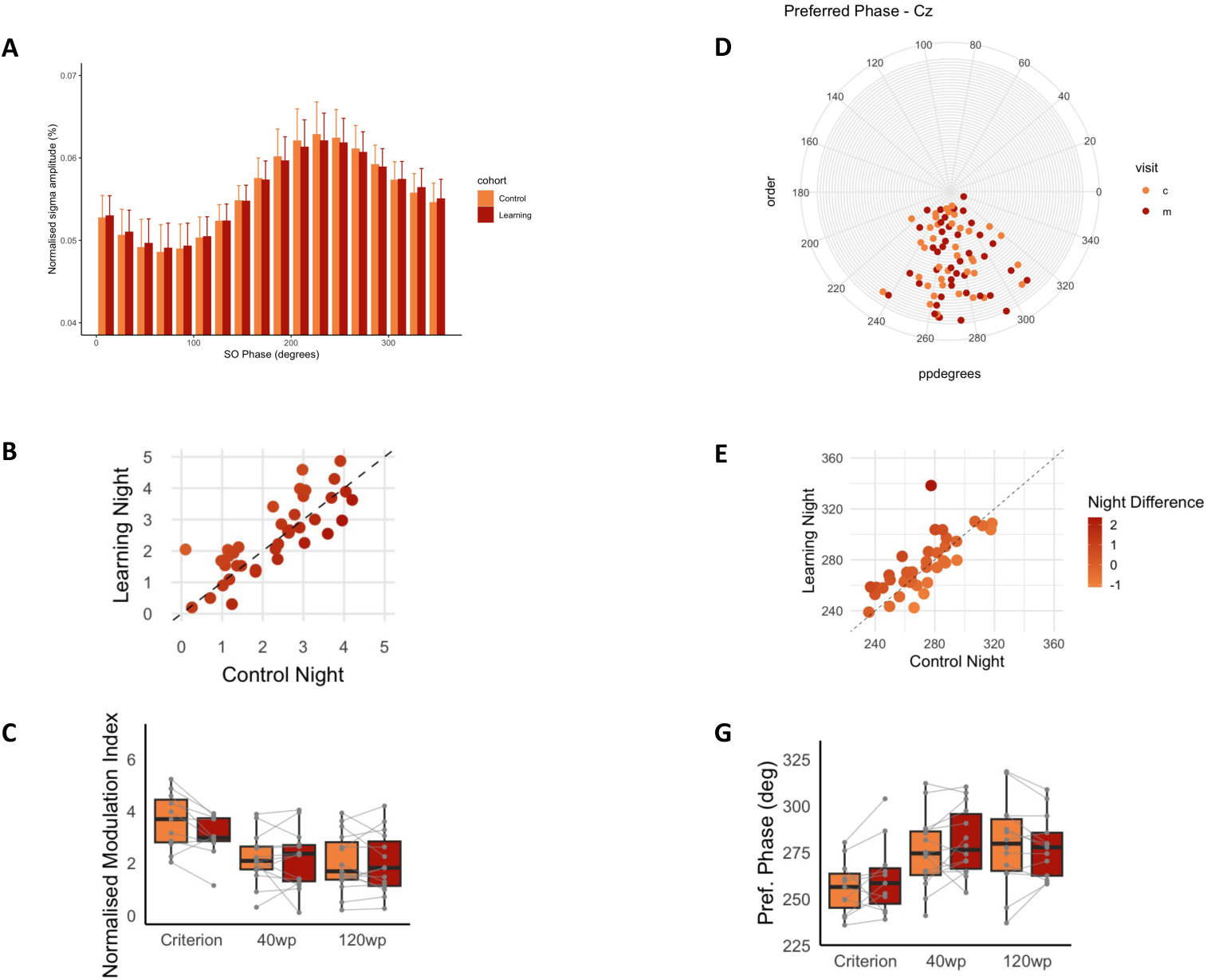
Measures of phase-amplitude coupling between slow oscillations (SO) and spindles on Cz during NREM sleep. **A**. Normalised amplitude of sigma frequencies distributed across 18 phase bins of the SO event for all participants. There were no night differences in amplitude within any phase bin. **B**. Between-night correlations of SO-spindle coupling strength measured by the modulation index. Light colors represent a greater number on the control night, darker colors represent a greater number on the learning night. **C**. There were no night differences in normalised strength (modulation index) of SO-spindle coupling per night across the 3 experiments. **D**. Circular plots showing the preferred phase of the maximal sigma amplitude per night. Each ring (distance from the centre) represents one participant. **E**. Between-night correlations of SO-spindle coupling strength measured by the modulation index. Light colors represent a greater number on the control night, darker colors represent a greater number on the learning night. **G**. There were no night differences in preferred coupling phase of SO-spindle coupling per night across the 3 experiments.

**Supplementary Table 1:**
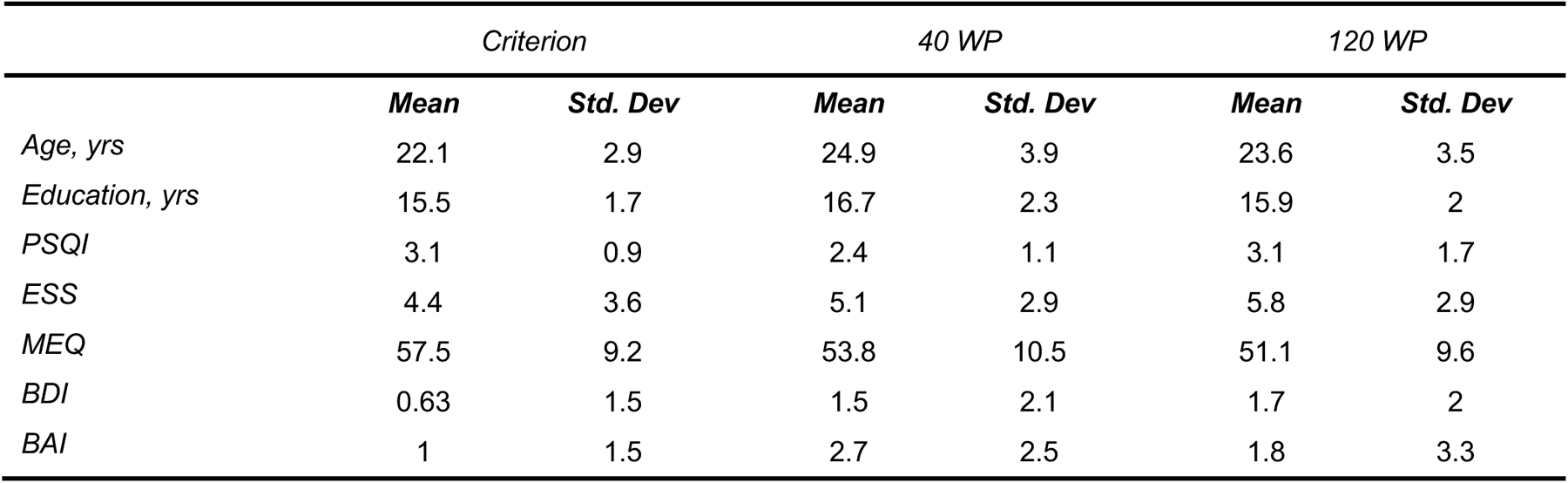
Demographics and scores on sleep and mood questionnaires.

**Supplementary Table 2:** Sleep architecture measures on Control and Learning nights.

| **Criterion** |  |  |  |  |  |  |  |
| --- | --- | --- | --- | --- | --- | --- | --- |
|  | **Control** |  | **Learning** |  | **t/W** | **p** | **g** |
|  | **Mean** | **Std. Dev** | **Mean** | **Std. Dev** |  |  |  |
| *TST* | 416.23 | 58.67 | 407.45 | 61.04 | 0.34 | 0.735 | 0.13 |
| *SL* | 11.23 | 10.54 | 10.09 | 5.82 | 31 | 0.894 | 0.12 |
| *SL_toREM* | 112.46 | 37.88 | 105.27 | 39.23 | 0.44 | 0.667 | 0.17 |
| *W_tsp* | 6.5 | 4.76 | 3.93 | 1.70 | 47 | 0.23 | 0.65 |
| *N1_tsp* | 3.46 | 1.91 | 3.15 | 1.49 | 0.42 | 0.683 | 0.16 |
| *N2_tsp* | 48.64 | 7.36 | 49.84 | 6.84 | -0.4 | 0.695 | 0.16 |
| *N3_tsp* | 19.96 | 6.0 | 21.22 | 5.34 | -0.52 | 0.611 | 0.20 |
| *REM_tsp* | 21.44 | 5.18 | 21.85 | 5.17 | 31 | 0.894 | 0.07 |
| **40 WP** |  |  |  |  |  |  |  |
|  | **Control** |  | **Learning** |  | **t/W** | **p** | **g** |
|  | **Mean** | **Std. Dev** | **Mean** | **Std. Dev** |  |  |  |
| *TST* | 394.9 | 85.86 | 399.2 | 52.48 | 66 | 0.755 | -0.06 |
| *SL* | 7.52 | 5.99 | 8.27 | 5.68 | 36 | 0.315 | -0.12 |
| *SL_toREM* | 71.69 | 13.62 | 94.23 | 54.12 | 21.5 | 0.056 | -0.55 |
| *W_tsp* | 5.26 | 3.52 | 5.47 | 7.25 | 76 | 0.379 | -0.03 |
| *N1_tsp* | 4.11 | 2.28 | 3.79 | 2.16 | 81 | 0.244 | 0.13 |
| *N2_tsp* | 40.84 | 5.87 | 44.95 | 9.49 | -1.43 | 0.167 | -0.49 |
| *N3_tsp* | 25.36 | 9.49 | 25.19 | 9.55 | 51 | 0.629 | 0.02 |
| *REM_tsp* | 24.44 | 5.56 | 20.6 | 5.7 | 1.87 | 0.072 | 0.64 |
| **120 WP** |  |  |  |  |  |  |  |
|  | **Control** |  | **Learning** |  | **t/W** | **p** | **g** |
|  | **Mean** | **Std. Dev** | **Mean** | **Std. Dev** |  |  |  |
| *TST* | 398.67 | 55.66 | 411.9 | 32.48 | -0.8 | 0.435 | -0.25 |
| *SL* | 11.01 | 8.36 | 12.08 | 8.05 | 45 | 0.41 | -0.12 |
| *SL_toREM* | 91.74 | 17.03 | 98.18 | 49.21 | 49 | 0.551 | -0.14 |
| *W_tsp* | 3.81 | 3.03 | 3.22 | 2.9 | 78 | 0.32 | 0.19 |
| *N1_tsp* | 6.94 | 3.96 | 6.94 | 3.85 | 60 | 0.999 | <0.01 |
| *N2_tsp* | 47.11 | 5.01 | 48.13 | 5.06 | -0.55 | 0.585 | -0.19 |
| *N3_tsp* | 22.47 | 6.22 | 22.18 | 7.72 | 0.11 | 0.91 | 0.04 |
| *REM_tsp* | 19.66 | 4.62 | 19.54 | 5.53 | 0.07 | 0.948 | 0.02 |

**Supplementary Table 3:** Sleep mico-architecture measures on Control and Learning nights.

| Criterion |  |  |  |  |  |  |  |
| --- | --- | --- | --- | --- | --- | --- | --- |
|  | Control |  | Learning |  | t/W | p | g |
|  | Mean | Std. Dev | Mean | Std. Dev |  |  |  |
| Spindles |  |  |  |  |  |  |  |
| Density (n•min) | 1.25 | 0.23 | 1.26 | 0.24 | -0.08 | 0.94 | -0.03 |
| Duration (sec) | 0.78 | 0.02 | 0.78 | 0.03 | 0.07 | 0.949 | 0.03 |
| Amplitude (μV²) | 257.63 | 57.07 | 260.74 | 55.66 | -0.13 | 0.898 | -0.05 |
| SOs |  |  |  |  |  |  |  |
| Density (n•min) | 3.86 | 0.75 | 4.09 | 0.74 | -1.44 | 0.179 | -0.28 |
| Duration (sec) | 1.34 | 0.07 | 1.32 | 0.08 | 0.4 | 0.69 | 0.15 |
| Amplitude (μV²) | 354.97 | 68.4 | 351.49 | 60.08 | 0.13 | 0.9 | 0.04 |
| 40 WP |  |  |  |  |  |  |  |
|  | Control |  | Learning |  | t/W | p | g |
|  | Mean | Std. Dev | Mean | Std. Dev |  |  |  |
| Spindles |  |  |  |  |  |  |  |
| Density (n•min) | 1.39 | 0.23 | 1.38 | 0.23 | 0.14 | 0.886 | 0.05 |
| Duration (sec) | 0.78 | 0.03 | 0.79 | 0.03 | -0.5 | 0.622 | -0.17 |
| Amplitude (μV²) | 131.24 | 36.88 | 131.93 | 36.24 | -0.05 | 0.959 | -0.02 |
| SOs |  |  |  |  |  |  |  |
| Density (n•min) | 4.69 | 1.09 | 4.5 | 0.96 | 81 | 0.244 | 0.18 |
| Duration (sec) | 1.26 | 0.09 | 1.26 | 0.07 | 0.2 | 0.843 | 0.07 |
| Amplitude (μV²) | 186.8 | 43.22 | 192.33 | 50.88 | -0.32 | 0.751 | -0.11 |
| 120 WP |  |  |  |  |  |  |  |
|  | Control |  | Learning |  | t/W | p | g |
|  | Mean | Std. Dev | Mean | Std. Dev |  |  |  |
| Spindles |  |  |  |  |  |  |  |
| Density (n•min) | 1.3 | 0.24 | 1.26 | 0.26 | 0.36 | 0.718 | 0.12 |
| Duration (sec) | 0.8 | 0.04 | 0.8 | 0.04 | -0.01 | 0.992 | 0 |
| Amplitude (μV²) | 153.05 | 44.07 | 157.38 | 43.66 | -0.27 | 0.789 | -0.09 |
| SOs |  |  |  |  |  |  |  |
| Density (n•min) | 3.39 | 0.54 | 3.43 | 0.51 | 47 | 0.478 | -0.08 |
| Duration (sec) | 1.37 | 0.05 | 1.36 | 0.05 | 0.06 | 0.951 | 0.02 |
| Amplitude (μV²) | 198.14 | 45.63 | 192.23 | 38.37 | 0.38 | 0.704 | 0.11 |

**Supplementary Table 4:** Prediction of word-pair consolidation by SO-sigma mean vector angle (MVA) and resultant vector length (RVL) from Fz and Pz during all NREM sleep on the learning night. N = 11.

| | *F* | $\Delta R^2$ | *p* | *B* (SE) | *eta*<sup>2</sup> |
| --- | --- | --- | --- | --- | --- |
| Fz |  |  |  |  |  |
| *Model* | 1.39 | 0.26 | 0.304 |  |  |
| *MVA* | 2.69 |  | 0.139 | -0.51 (0.31) | 0.25 |
| *RVL* | 0.01 |  | 0.948 | 0.02 (0.30) | 0.01 |
| Pz |  |  |  |  |  |
| *Model* | 3.6 | 0.47 | 0.077 |  |  |
| *MVA* | 3.68 |  | 0.091 | 0.30 (0.29) | 0.24 |
| *RVL* | 0.61 |  | 0.459 | -0.22 (0.29) | 0.03 |

